# Impaired Fermentation in the Absence of Gut Microbiota Promotes Inulin-Induced Intestinal Inflammation

**DOI:** 10.64898/2026.11.06.764925

**Authors:** Edward Moncada, Paola Andino, Vignesh Nathan, Yunus E. Tunçil, Stephen R. Lindemann, Beng San Yeoh, Andrew T. Gewirtz, Tzu-Wen L. Cross, Matam Vijay-Kumar, Lavanya Reddivari

## Abstract

**Background and aims:** Inflammatory bowel diseases (IBD) are characterized by altered gut microbiota composition and impaired dietary fiber fermentation, which can complicate symptom management. Although inulin is a well-established prebiotic fiber, it can exacerbate gut inflammation under certain conditions. We hypothesized that impaired microbial fermentation leaves intact inulin in the gut to activate epithelial immune receptors and promote inflammation, whereas effective fermentation reduces this pro-inflammatory signaling.

**Methods:** We used germ-free (GF) and conventional (CONV) interleukin-10 knockout (Il10⁻/⁻) and wild-type (WT) C57BL/6J mice to assess the role of microbial fermentation and host genetic susceptibility in IBD. GF Il10⁻/⁻ mice were colonized with fecal microbiota from ulcerative colitis (UC) patients or healthy individuals with fiber intolerance (FI). Mice were fed inulin or structurally distinct, physicochemically matched fiber analogs. Colonic inflammatory and barrier-related gene expression and protein levels were assessed by qPCR and ELISA.

**Results:** In GF Il10⁻/⁻ mice, inulin, but not cellulose, induced weight loss, diarrhea, and rectal prolapse. Increased colonic toll-like receptor expression (TLR2, TLR4) as well as elevated serum amyloid A (SAA) and lipocalin-2 levels indicated immune activation by unfermented inulin in GF mice. Colonization with dysbiotic microbiota attenuated these effects. β-fructan fibers including fructooligosaccharides and chicory-derived inulin elicited marked inflammatory responses, whereas agave-derived inulin did not, highlighting the influence of fiber composition in inflammation associated with IBD.

**Conclusions:** Effective microbial fermentation is essential for host tolerance to fermentable fibers. These findings underscore the need of personalized nutrition when recommending fiber supplementation in immune-compromised individuals or those with gut dysbiosis.

**Summary:** This study shows that microbial fermentation is essential for host tolerance to dietary fibers, and that fiber structure determines the immunological response.

**Key Messages:** *What is already known?:*

- Dietary fiber effects are not universally beneficial; for example, inulin, a commonly used prebiotic, has been shown to exacerbate colitis symptoms in animal models.

*What is new here?:*

- Inulin induced severe inflammation in germ-free Il10⁻/⁻ mice, but this phenotype was absent in conventionally raised mice with the same genetic predisposition.
- Even dysbiotic microbial colonization attenuated inulin’s negative physiological effects.
- Not all fructans behave equally; chicory inulin and fructooligosaccharides (FOS) elicited pronounced inflammatory responses, whereas agave inulin did not.

*How can this study help patient care?:*

- These findings suggest that when selecting dietary fiber for individuals with IBD, microbial fermentative capacity and fiber properties should be considered.

## 1. Introduction

Dietary fibers are a class of plant-derived complex carbohydrates resistant to host enzymatic digestion in the upper gastrointestinal tract, thus reaching the distal part of the intestine in intact form^1^. In the colon, undigested fibers are metabolized via fermentation by gut-resident microbes predominantly into short-chain fatty acids (SCFA), which can substantially influence host physiology. Among these fibers, inulin, a β (2→1)-linked fructan composed of linear chains of fructose with a terminal glucose unit^2^, is widely studied for its prebiotic and bifidogenic potential, stimulating the growth of beneficial bacteria^3^. Through fermentation of inulin, the gut bacteria produce SCFAs, principally acetate, propionate, and butyrate, in mM levels, which have been shown to be largely anti-inflammatory and reinforce mucosal epithelial barrier integrity^4,5^. Beyond SCFAs, however, inulins are fermented to a multitude of metabolites in ways that depend on inulin structure; these microbial metabolites may also be influential on health^6^.

Microbial fermentation is central to the health-promoting actions of inulin. SCFAs derived from fermentation (especially butyrate) serve as energy sources for colonocytes and modulate immune cell differentiation, including the induction of regulatory T cells (Tregs)^7^. In GF mice, which lack microbial fermentation capacity, accumulation of unfermented inulin has been associated with epithelial stress and immune activation^8^, suggesting that microbial metabolism is essential to obtain fiber health benefits^9^. Host-specific responses to inulin sensitivity have been intensively linked to gut microbiota composition^10,11^, since this dictates fermentability of inulin and the immune outcomes. Microbial profiles having higher abundances of *Bacteroides* or *Prevotella* may exhibit distinct fermentation patterns and SCFA profiles^12^ in response to inulin compared to microbial profiles with lower abundance. Metagenomic and metabolomic studies have shown that microbial metabolic capacity, not just taxonomic composition, dictates SCFA output and pro- and anti-inflammatory effects^13,14^.

However, accumulating evidence indicates that dietary inulin effects are not universally beneficial and may be context-dependent^15^. Although inulin typically promotes eubiosis and mucosal homeostasis, certain host and microbial profiles, such as those found in UC patients, can result in detrimental outcomes. In genetically susceptible animal models of IBD or individuals with gut dysbiosis, inulin has been linked to exacerbation of intestinal inflammation, hepatotoxicity, and even cholestatic liver cancer^16^. Recent studies have shown that microbiota-accessible carbohydrates, including inulin, can provoke intestinal inflammation in a manner that is microbiota-specific^17^.

Host genetics are also a critical determinant in shaping the physiologic and immunologic outcomes of dietary fiber consumption. One well-established model for studying genetic susceptibility to intestinal inflammation is the Il10^⁻/⁻^ mouse model, which lacks the regulatory anti-inflammatory cytokine IL-10 and consequently develops spontaneous colitis due to unrestrained immune activation in response to microbial and dietary antigens^18^. These mice provide a robust platform for assessing the pro-or anti-inflammatory potential of specific dietary components under conditions of compromised immune response. This model allows us to assess how deficiencies in key anti-inflammatory pathways regulate host immune response to fiber supplementation; such insight is critical for understanding how the immune dysregulation or other immune-mediated conditions govern dietary outcomes, paving the way for personalized rather than one-size-fits-all fiber interventions^15^.

The structural properties of dietary fibers such as solubility, chain length, fermentation rate, monomeric unit composition, linkage types and patterns also govern their influence on fermentability and immunogenic potential^19,20^. For instance, dietary fibers with a lower degree of polymerization may be more rapidly fermented in the proximal colon, whereas long-chain variants persist longer and may be fermented by distinct microbial populations in the distal colon^21^. These kinetic and spatial differences in fermentation can influence SCFA production, bioavailability, and whether inflammatory signaling is activated or suppressed^22,23^. Comparative studies have shown that even subtle changes in fiber structure can elicit divergent microbial and immunological responses, emphasizing the need for precise fiber characterization for personalized nutrition^15,25^.

Thus, although inulin is often considered beneficial due to its prebiotic and SCFA-generating properties, its effects are highly dependent upon host-microbe interactions in the gut, immune status, and fiber structure. The current study aims to evaluate these variables to understand the mechanisms through which certain fibers exacerbate or ameliorate inflammation. We found that, in the absence of microbial fermentation, intact inulin polymers interact with colonic Toll-like receptors and amplify both gut and systemic inflammation in a genetically susceptible host, whereas these effects can be prevented by the presence of conventional and functional gut microbiota. Even partial bacterial colonization from dysbiotic donors was sufficient to attenuate these responses, indicating that microbial metabolic activity, rather than host genetics, is the primary determinant of fiber tolerance. Importantly, not all fibers behave equally; chicory-derived inulin and fructooligosaccharide (FOS) drove pronounced inflammatory responses, while agave-derived inulin did not, implicating fiber-specific structural and chemical properties as key modulators of intestinal inflammation. Taken together, our study contributes to the precise fiber selection for clinical and nutritional applications, particularly for vulnerable populations such as those with IBD or gut microbial dysbiosis.

## 2. Materials and Methods

### 2.1 Consent and Ethics Approval

Consent was obtained from patients/guardians, and the protocol for sample collection was approved by Purdue’s Institutional Review Board (IRB-2021-109). Additionally, all animal procedures were approved by the Purdue Institutional Animal Care and Use Committee (PIACUC approval No. 1810001817) according to the recommendations of the Guide for the Care and Use of Laboratory Animals published by the National Institutes of Health.

### 2.2 Human Donor Criteria and Sample Collection

Patients with confirmed ulcerative colitis (UC) (fecal calprotectin levels above 1,900 μg/g) and fiber intolerance (FI) conditions were used as donors for mice colonization and *in vitro* fecal fermentation. FI donor status was confirmed through a questionnaire that determined sensitivity to high-fiber foods. Fecal samples were collected using sterile fecal collection kits, subsequently aliquoted, and processed under aseptic and anaerobic conditions. Samples were then stored at - 80°C and used within a month of freezing to preserve microbial integrity for subsequent transplantation into recipient mice and application in *in vitro* fecal fermentation assays.

### 2.3 Dietary Fibers and Experimental Design

Dietary fibers were selected based on their distinct structural and physicochemical characteristics. As previously reported by our laboratory^26^, host responses to dietary fiber supplementation are influenced by both fiber-specific and host-dependent variables. Physicochemical properties of fibers that are considered potential modulators of inflammatory processes are listed in Table S1. This study focuses particularly on inulin, where its potentially detrimental effects on gut health in certain individuals remain poorly understood^8^.

Male and female mice aged five to six weeks (n = 6 - 10) were randomly allocated to different fiber diets for two weeks (Fig. S1). Throughout the intervention period, animals were monitored daily for potential adverse effects, including changes in body weight, stool consistency, and food consumption. Upon completion of the dietary intervention, mice were euthanized, and post-necropsy analyses were conducted to assess treatment-related outcomes.

### 2.4 Mouse Models and Diets

All diets utilized in this study were manufactured by Envigo (Madison, WI, USA). GF mice were bred in the dedicated Gnotobiotic Animal Facility at Purdue University. After weaning, the mice were kept on the sterile irradiated AIN-93G diet. Five to six weeks old mice were taken out from the breeding isolators and maintained in the isocages for the entire experimental period. Conventionally raised specific pathogen free mice (CONV) were bred and maintained in Purdue University’s SPF Mouse Facility. Mice were housed under 12:12-h light-dark cycles, controlled temperature (20–25 ◦C) and relative humidity (40–60%). Experimental diets were formulated to contain 15% (w/w) of fiber (Table S3).

### 2.5 Fecal Microbiota Transplantation

Frozen human fecal samples were homogenized in pre-reduced sterile phosphate-buffered saline (PBS) at a ratio of 1.0 g feces to 3 mL PBS to prepare a fecal slurry. The slurry was centrifuged at 2,000 x g for 10 minutes at 4 °C, and the resulting supernatant was collected. Mice were administered 150 μL of fecal supernatant via oral gavage twice during the first week, with a 72-hour interval between each administration^26^.

### 2.6 Mouse Necropsy and Sample Processing

After the experimental period, mice were euthanized via carbon dioxide asphyxiation. Whole blood was collected by cardiac puncture, transferred into capillary collection tubes to isolate serum, which was subsequently stored at −80 °C for future analyses. Following the necropsy, organs and gastrointestinal tissues were excised. Luminal contents were gently flushed, and tissues were rinsed with sterile PBS. Pre- and post-digesta tissue weights were recorded. The cecum was flash-frozen in dry ice, while the colon was longitudinally opened and sectioned into proximal and distal regions. Each section was further subdivided for downstream applications, including RNA extraction for gene expression analysis, snap-frozen for protein assays, and fixed in 10% neutral-buffered formalin and Carnoy’s solution for histological evaluation and mucus architecture, respectively.

### 2.7 RNA Isolation and Gene Expression Analyses

Total RNA was extracted from frozen colon tissue samples using the PureLink RNA Mini Kit (Invitrogen, Carlsbad, CA, USA) following the manufacturer’s protocol. RNA concentrations were quantified using the BioTek Cytation 3 Cell Imaging Multi-Mode Reader (Thermo Fisher Scientific Inc., Winooski, VT, USA). Complementary DNA (cDNA) synthesis was performed using the qScript cDNA SuperMix (Quantabio, Beverly, MA, USA) following the manufacturer’s guidelines. Quantitative real-time PCR (qRT-PCR) was conducted on a Roche LightCycler 96 instrument using PerfeCTa® SYBR® Green FastMix (Quantabio, Beverly, MA, USA). The thermal cycling conditions included an initial denaturation at 95 °C for 10 minutes, followed by 45 amplification cycles consisting of 95 °C for 10 seconds (denaturation) and 55 °C for 60 seconds (annealing/extension). Gene expression levels were normalized to β-actin as the internal reference gene^27^. Primer sequences for the analyzed genes are listed in Supplementary Table S2.

### 2.8 Colon Histology

Two distal colon sections were collected from each mouse for histological analysis. The first section was fixed in 10% neutral buffered formalin at room temperature for a minimum of 24 hours, paraffin-embedded, and sectioned at 4 µm. The second section was fixed in Carnoy’s solution at room temperature for a minimum of 24 hours, paraffin-embedded, and sectioned at 4 µm, then stained with Alcian blue for mucus-specific assessment. All slides were examined by light microscopy for histological evaluation and mucus layer thickness measurements.

### 2.9 Lipocalin-2 and Serum Amyloid A quantification in fecal and serum samples

Serum amyloid A (SAA), lipocalin-2 (Lcn2) and fecal Lcn2 levels were measured using a DuoSet ELISA kit (R&D Systems, Minneapolis, MN, USA) according to the manufacturer’s instructions. Briefly, samples and standards were added to antibody-coated plates, followed by incubation with detection antibody and streptavidin-horseradish peroxidase. Signal development was achieved using a chromogenic substrate and absorbance was measured at 450 nm. Concentrations were calculated from a standard curve, and samples were analyzed in triplicate.

### 2.10 Fructose Quantification in Cecal Digesta

To measure the extent of inulin degradation, fructose levels were quantified in fecal and cecal digesta using high-performance liquid chromatography (HPLC) analysis. Briefly, samples were weighed and diluted in deionized water at appropriate ratios (e.g., 1:2–1:10, depending on sample consistency and expected sugar concentrations), followed by thorough homogenization via vortexing. Homogenates were centrifuged (700 x g for 2 min), and the supernatant was collected and mixed with an equal volume of 5.0 M trifluoroacetic acid (TFA). Samples were then subjected to acid hydrolysis at 121 °C for 90 min. Following hydrolysis, TFA was evaporated to dryness, and residues were washed twice with 100% methanol with intermediate drying to ensure complete removal of residual TFA. Dried samples were subsequently reconstituted in deionized water, filtered through 0.22 μm syringe filters, and transferred to vials for analysis. Quantification of simple sugars was performed using HPLC before and after acid hydrolysis. Final volumes were adjusted according to initial sample mass and dilution factors.

### 2.11 Statistical Analysis

Data were analyzed and graphed in R software and GraphPad prism 10. Outliers were identified using the robust regression and outlier removal (ROUT) method. Data sets were analyzed for normality using the Shapiro-Wilk test, and if variables passed the test, then they were analyzed utilizing t-test, one-way, and two-way ANOVA. For variables not following normality, a Mann–Whitney or a Kruskal–Wallis test followed by Dunn’s multiple comparison test was used. Data are presented as the mean ± standard error of the mean (SEM). The p-values below 0.05 were considered statistically significant. Statistical significance is indicated as follows: ns = no significance, * = p < 0.05; ** = p < 0.01; *** = p < 0.001, **** = p < 0.0001.

## 3. Results

### 3.1 Unfermented dietary chicory inulin, but not unfermented cellulose, triggers inflammatory responses in germ free Il10^-/-^ mice

Previous *ex vivo* and *in vitro* studies have demonstrated that unfermented fructans, such as inulin, may elicit pro-inflammatory responses in a subset of individuals with IBD^8^. To assess whether the absence of gut microbial fermentation contributes to such effects *in vivo*, we utilized GF and CONV Il10^⁻/⁻^ mouse models and administered fiber diets containing either cellulose or inulin (15% w/w). As shown in Figure 1A–B, mice fed cellulose exhibited no significant changes in body weight in either GF or CONV conditions (Fig. 1A). No signs of inflammation or distress were observed. In contrast, GF Il10^⁻/⁻^ mice receiving inulin exhibited a progressive and significant reduction in body weight, exceeding 20% (IACUC guided mortality body weight loss) by the end of the experimental period (Fig. 1B). This was accompanied by clinical signs of gastrointestinal distress including persistent diarrhea (Fig. 1D) and rectal prolapse, a severe complication of colitis (Fig. 1E). CONV Il10^⁻/⁻^ mice supplemented with inulin and with intact gut microbiota maintained stable body weight and displayed no pathological symptoms or adverse effects, unlike the GF Il10- /- mice (Fig. 1B).

**Fig 1.**
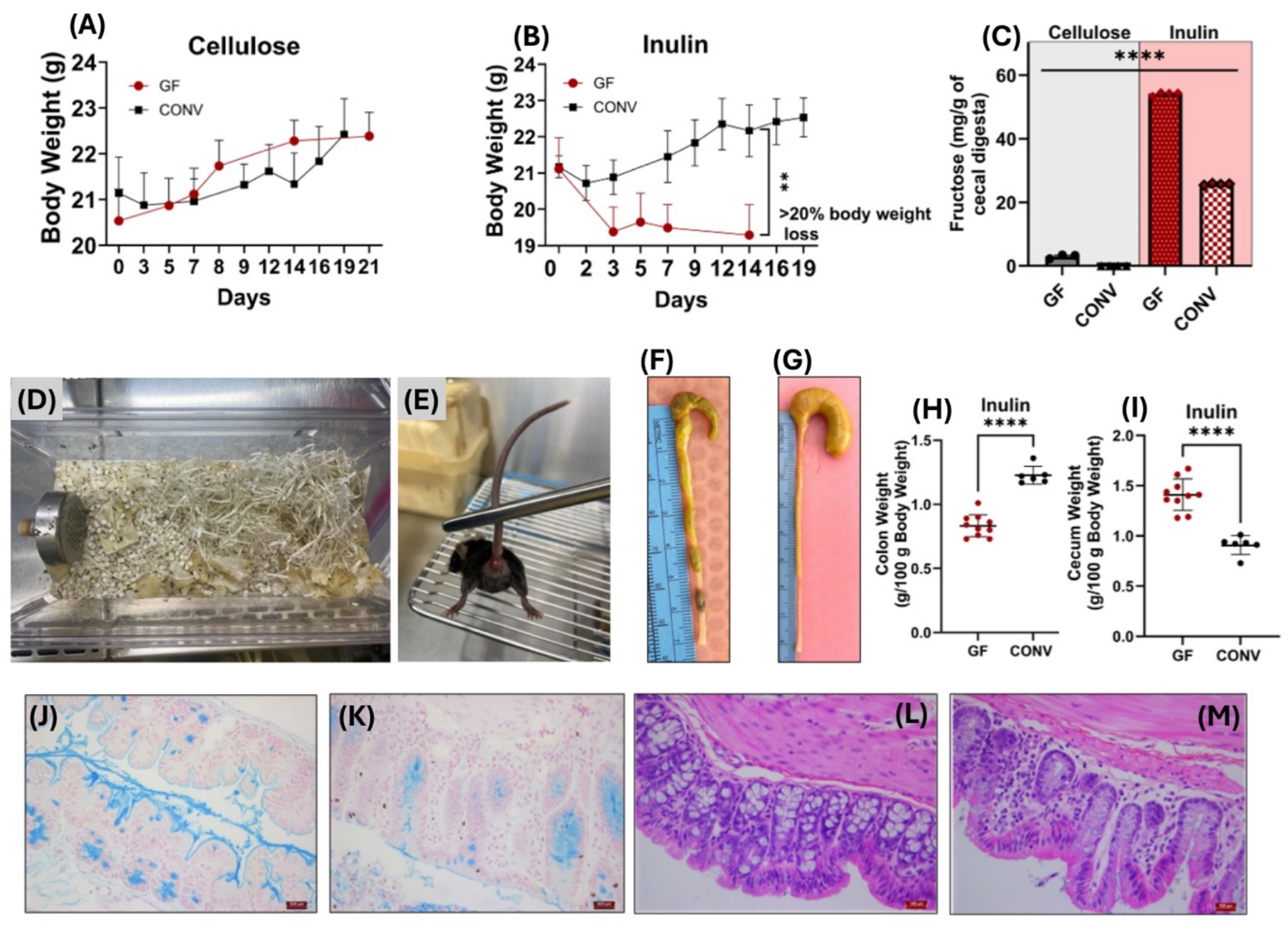
Mice (Il10^⁻/⁻^) body weights throughout the experimental period for cellulose (A) and inulin (B) treatment groups in GF and CONV conditions. Fructose concentrations in cecal digesta from GF and CONV mice fed diets containing either cellulose or inulin, used as an indicator of residual (unfermented) inulin within the cecal contents (C). Cage bedding showing mouse diarrhea severity (D) and rectal irritation/prolapse (E) in germ-free mice fed with inulin. Cecum and colon from Il10^⁻/⁻^ mice supplemented with inulin having their native microbiota (F) and GF (G). Colon (H) and cecum (I) weights from inulin-fed GF and CONV mice. Representative Alcian blue-stained distal colon sections from GF Il10⁻/⁻ mice fed cellulose (J) or inulin (K). Cellulose-fed mice (J) displayed abundant and continuous mucus staining, indicative of intact goblet cell populations and a well-preserved mucus layer. In contrast, inulin-fed mice (K) exhibited markedly reduced Alcian blue staining intensity and goblet cell content, suggesting mucus layer depletion and compromised epithelial barrier integrity. Representative H&E-stained distal colon sections from GF Il10⁻/⁻ mice fed cellulose (L) or inulin (M), showed preserved crypt architecture and epithelial organization in cellulose-fed mice (L), whereas inulin-fed mice (M) exhibited altered crypt morphology and increased cellular infiltration within the lamina propria, consistent with intestinal tissue inflammation. Scale bars = 500 µm. Replicates (n = 6–10) for each mice group are plotted. Asterisks (**) and horizontal lines represent a statistical difference (p ≤ 0.01) between the two groups. (****) represents p ≤ 0.0001 and (ns) means no statistical difference. Data is shown as mean ± SEM. Data is shown as mean ± SEM. Alt text: Multi-panel figure showing body weights (A–B), fructose concentrations (C), images of cage bedding showing diarrhea and rectal prolapse (D–E), cecum and colon images (F–G), colon and cecum weights (H–I), Alcian blue-stained distal colon sections (J–K), and H&E-stained distal colon sections (L–M) in GF and CONV Il10⁻/⁻ mice fed cellulose- or inulin-containing diets.

To assess the presence of intact inulin, cecal digesta samples were hydrolyzed, and the resulting fructose concentration was quantified as a proxy for residual inulin (Fig. 1C). As expected, cellulose-fed mice exhibited negligible levels of fructose in the digesta. In contrast, GF mice fed inulin displayed significantly higher fructose concentrations compared to their CONV counterparts, indicating the lack of microbial fermentation and increased persistence of intact inulin in the absence of microbiota. These findings suggest that unfermented inulin may directly interact with the host gut epithelia, potentially contributing to the observed phenotypes, including weight loss, diarrhea, and colonic mucosal thinning.

General morphological analysis further revealed marked differences between the two models. Inulin-fed CONV mice had normal stool consistency and preserved colon morphology (Fig. 1F), whereas their GF counterparts displayed enlarged ceca and the absence of well-formed fecal pellets (Fig. 1G), indicative of unfermented inulin buildup due to the lack of gut microbial metabolism. Additionally, colonic thinning was observed in GF mice, which may reflect epithelial atrophy or compromised mucosal integrity secondary to unfermented fiber-induced osmotic stress and epithelial inflammation.

Post-necropsy organ weights were assessed to identify potential physiological markers of inflammation such as splenomegaly, colonic thickening or shortening, and cecal distension between the different mouse models. Colon length remained unaffected by dietary fiber supplementation in both GF and CONV Il10^⁻/⁻^ mice (Fig. S2A–B). However, colon weights were significantly higher in the CONV mice for both dietary groups (Fig. 1H and Fig. S2C). Consequently, the colon weight-to-length ratio differed significantly between models, with the most noticeable difference in the inulin-supplemented group (Fig. S2D–E), potentially indicating increased water retention or mucosal edema due to osmotic effects from unfermented inulin in GF mice. As expected, cecal weights were substantially elevated in the GF animals regardless of dietary treatment (Fig. 1I and Fig. S2F), consistent with the lack of microbial fermentation and consequent fiber accumulation in the cecum. Liver and spleen weights (Fig. S2) were not significantly altered by either fiber supplementation or microbial status, suggesting no hepatosplenomegaly or systemic immune activation in these models. In contrast, kidney weights were significantly elevated in the GF mice receiving inulin (Fig. S2), which may indicate renal stress or early signs of inflammation-associated fluid dysregulation. Histological examination of Alcian blue-stained distal colon sections provided further morphological evidence of mucosal compromise in inulin-fed GF mice. Cellulose-fed mice displayed abundant and continuous Alcian blue staining along the colonic epithelium, reflecting intact goblet cell populations and a well-preserved mucus layer (Fig. 1J). In contrast, inulin-fed GF mice exhibited markedly reduced staining intensity and goblet cell content (Fig. 1K), indicative of mucus layer depletion, and impaired epithelial barrier integrity. These histological findings are consistent with the observed colonic thinning and body weight loss; collectively suggesting that unfermented inulin disrupts mucus production and compromises the protective mucosal barrier.

To evaluate the potential for unfermented fiber to induce colonic inflammation, relative gene expression of colonic pattern recognition receptors, Toll-like receptors 2 and 4 (TLR2 and TLR4), were measured. These receptors play a central role in the innate immune response by recognizing microbial- and damage-associated molecular patterns and subsequently activating pro-inflammatory signaling cascades, including NF-KB activation and the induction of cytokines such as IL-1β, IL-6, and TNF-α^28,29^. Cellulose supplementation did not significantly alter TLR2 or TLR4 expression in GF mice compared to CONV mice (Fig. 2A-B). In contrast, GF mice supplemented with inulin exhibited a significant upregulation of both TLR2 and TLR4 expression relative to their CONV counterparts (Fig. 2A-B), indicating that unfermented inulin but not cellulose triggers pro-inflammatory responses. Additionally, systemic inflammatory markers were evaluated. Serum amyloid A (SAA), an acute-phase protein produced primarily by the liver in response to pro-inflammatory cytokines and widely used as an indicator of systemic inflammation in IBD, was significantly upregulated in inulin-fed GF mice (Fig. 2C). Similarly, lipocalin-2 (LCN-2), an acute phase protein that is strongly induced during intestinal inflammation and commonly used as a sensitive biomarker of intestinal inflammation in murine models of colitis, was measured in both fecal and serum samples. In feces, LCN-2 levels were significantly elevated in GF mice across both cellulose- and inulin-fed groups. In contrast, a significant increase in serum LCN-2 was observed only in inulin-fed mice. This pattern suggests that while the absence of gut microbiota promotes a baseline elevation in intestinal inflammation, as reflected by fecal LCN-2, systemic inflammatory responses are further exacerbated specifically by dietary inulin in GF conditions, indicating a dietary fiber-dependent exacerbation of inflammation beyond the gut.

**Fig 2.**
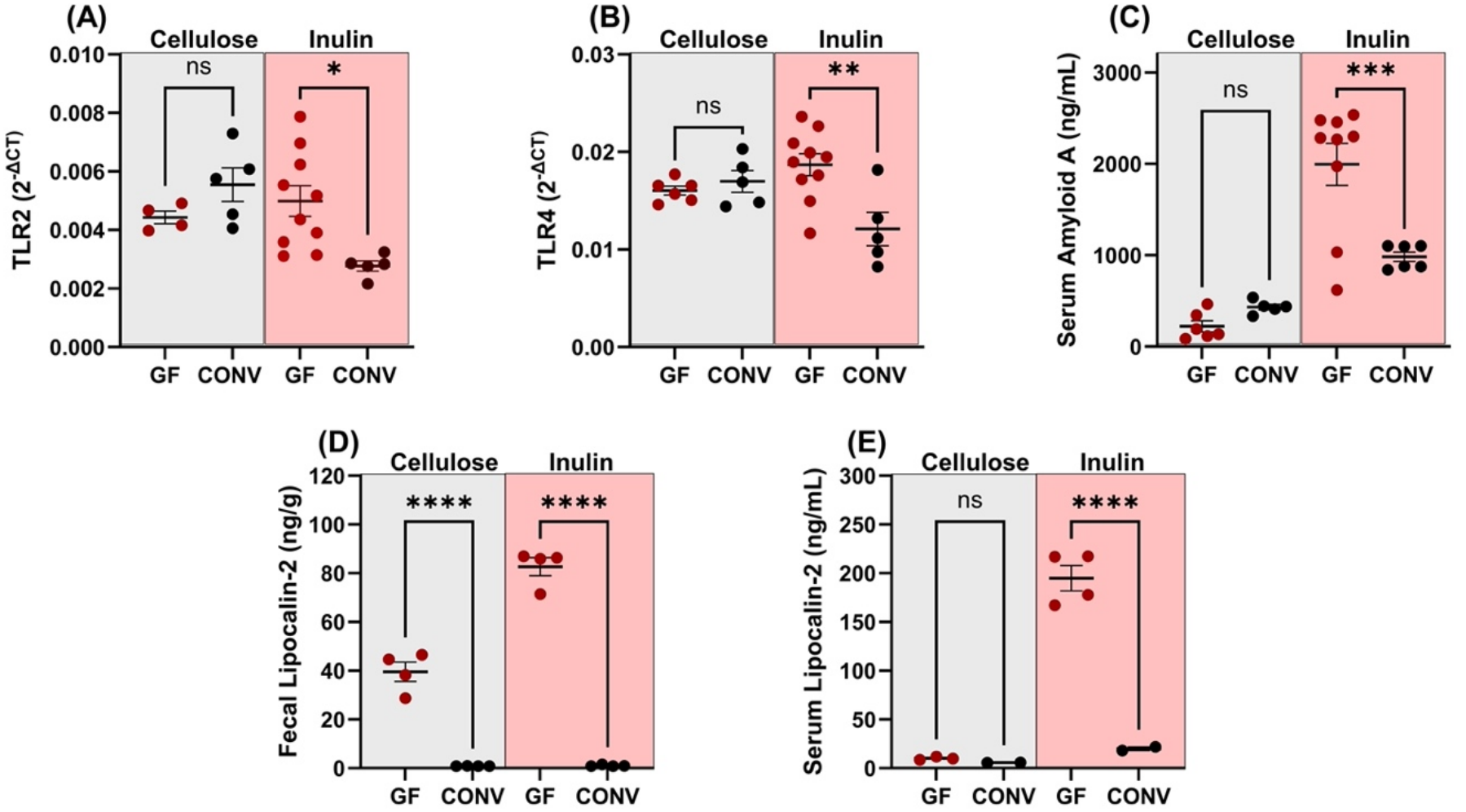
Relative gene expression of Toll-like receptors 2 (A) and 4 (B) in distal colon tissues of mice fed cellulose-or inulin-containing diets. Serum amyloid A concentrations (C), fecal lipocalin-2 concentrations (D), and serum lipocalin-2 concentrations (E) are also shown. Replicates (n = 4–10) for each mice group are plotted. An asterisk (*) and horizontal line represent a statistical difference (p ≤ 0.05) between the two groups. (**) represents p ≤ 0.01, (**) represents p ≤ 0.001, (**) represents p ≤ 0.0001 and (ns) means no statistical difference. Data is shown as mean ± SEM. Alt text: Multi-panel figure showing relative Toll-like receptor 2 (A) and 4 (B) gene expression in distal colon tissues, serum amyloid A concentrations (C), fecal lipocalin-2 concentrations (D), and serum lipocalin-2 concentrations (E) in mice fed cellulose-or inulin-containing diets.

### 3.2 Bacterial colonization from distinct donors reduced inulin associated inflammation in Il10^-/-^ mice

Given that unfermented inulin induces inflammation in GF mice, we next investigated whether microbiota transplantation from a donor with UC and an identified FI individual could modulate host responses to fiber supplementation. As shown in Figure 3A, cellulose-fed mice exhibited a net increase in body weight over the two-week supplementation period, with FI-recipient mice displaying a significant gain exceeding 10% of their baseline weight by week two. In contrast, all inulin-fed mice, regardless of microbial status, experienced body weight loss after one week of supplementation (Fig. 3B). Notably, by week two, mice colonized with either UC or FI microbiota showed partial recovery of body weight, while GF mice continued to lose weight. This temporal pattern suggests that as bacterial communities engraft and begin to function metabolically, the severity of weight loss in response to dietary inulin is attenuated.

**Fig 3.**
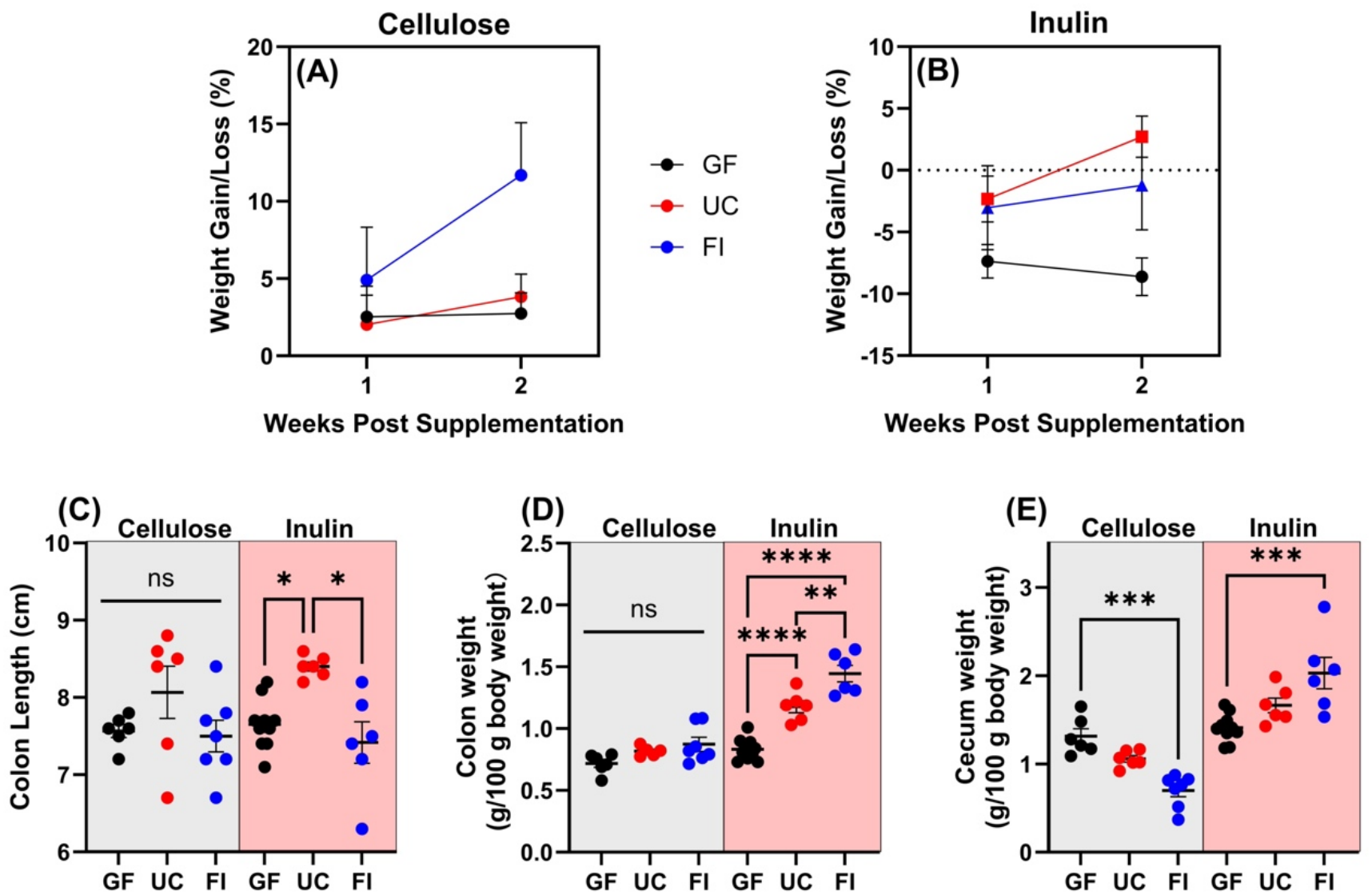
Mouse body weight gain or loss percentage after one-or two-week supplementation with cellulose (A) or inulin (B) under GF conditions, colonized with bacteria from a patient diagnosed with UC or FI. Mice colon lengths (C) and organ weights (D-E) for cellulose and inulin treatments in GF, and UC and FI colonized conditions. Replicates (n = 6–10) for each mice group are plotted. An asterisk (*) and horizontal line represent a statistical difference (p ≤ 0.05) between the two groups. (**) represents p ≤ 0.01, (**) represents p ≤ 0.001, (**) represents p ≤ 0.0001 and (ns) means no statistical difference. Data is shown as mean ± SEM. Alt text: Multi-panel figure showing mouse body weight gain or loss percentages after cellulose (A) and inulin (B) supplementation, colon lengths (C), colon weights (D), and cecum weights (E) in GF mice and mice colonized with microbiota from UC or FI donors.

Post-necropsy analysis of organs revealed that colon length remained unaffected by cellulose supplementation across all colonization models (Fig. 3C). In contrast, inulin-fed mice in the GF and FI groups exhibited significantly shorter colons compared to the UC-transplanted group (Fig. 3C). Colon weights were unchanged by cellulose fed group (Fig. 3D) but were elevated in the UC and FI groups following inulin supplementation (Fig. 3D). Cecum weight was significantly reduced in the FI-colonized mice receiving cellulose (Fig. 3E), whereas inulin-fed mice showed significant differences only between the GF and FI groups (Fig. 3E). Overall, these organ weights were largely unaffected by cellulose supplementation across colonization conditions. In contrast, inulin feeding elicited distinct, microbiota-dependent responses, which highlights the diverse physiological impact of dietary inulin compared to cellulose. Liver, spleen, and kidney weights exhibited minimal changes with inulin supplemented diet , whereas cellulose induced selective, microbiota-dependent effects (Fig. S3).

To evaluate the effects of microbial colonization on dietary fiber-induced inflammatory responses and colonic barrier function, relative gene expression of inflammatory markers (TLR2, TLR4, IL-6), fecal and serum levels of SAA and LCN-2, the epithelial tight junction protein Claudin-1, and the mucin-secreting gene Muc-2, were assessed. Colonization with microbiota derived from UC or FI donors resulted in reduced colonic expression of IL-6 in cellulose-fed mice (Fig. 4A), whereas no significant differences were observed among inulin-fed groups. The elevated IL-6 expression in GF cellulose-fed mice may reflect a transient basal immune activation in the absence of gut microbiota, as IL-6 is a rapidly inducible and short-lived cytokine. In contrast, the lack of difference in the inulin-fed groups may indicate that inulin-driven inflammatory responses involve alternative downstream pathways that do not prominently rely on sustained IL-6 expression. Furthermore, microbial colonization did not significantly alter TLR2 expression compared to GF controls (Fig. 4B). In contrast, TLR4 expression was significantly reduced in mice colonized with either UC- or FI-associated microbiota, a trend consistent across both dietary cellulose and inulin fed groups (Fig. 4C), suggesting a microbiota-mediated modulation of innate immune signaling. SAA levels revealed opposing trends between fiber types. In cellulose-fed mice, colonization with UC- or FI-associated microbiota increased SAA levels relative to GF controls. Conversely, in the inulin-fed groups, GF mice exhibited significantly higher SAA levels compared to colonized counterparts (Fig. 4D), indicating heightened systemic inflammation in the absence of gut microbiota under inulin diet fed groups. Assessment of localized inflammation using fecal LCN-2 showed that colonization with microbiota from either donor significantly reduced LCN-2 levels across both fiber treatments. Notably, GF mice fed inulin exhibited the highest fecal LCN-2 levels among all groups (Fig. 4E), indicating exacerbated intestinal inflammation in the absence of microbial fermentation. Consistent with this, serum LCN-2 concentrations were elevated in all inulin-fed groups relative to cellulose-fed treatments, with no significant differences among colonization conditions. The levels of circulating LCN-2 (∼150–200 ng/mL) in the inulin-fed groups are indicative of a pronounced systemic inflammatory state (Fig. 4F). With respect to gut barrier function, Claudin-1 expression was not significantly affected by microbial colonization in either dietary group (Fig. 4G). In contrast, Muc-2 expression was modulated in a microbiota colonization-dependent manner. In cellulose-fed mice, UC-derived microbiota colonization increased Muc-2 expression, whereas FI colonization significantly reduced its expression relative to both GF and UC groups (Fig. 4H). A similar pattern was observed under inulin fed group, with FI-colonized mice consistently exhibiting reduced Muc-2 expression. This reduction may reflect impaired mucus layer production associated with FI-derived microbiota, potentially compromising barrier integrity and contributing to increased susceptibility to inflammation. Bacterial colonization, even from dysbiotic sources such as UC and FI donors, can partially ameliorate the negative physiological effects associated with inulin supplementation, particularly the weight loss observed in GF mice. This attenuation appears to be associated with early-stage bacterial engraftment and partial restoration of fiber metabolism. Bacterial presence and fiber metabolism significantly reduced inflammatory markers such as IL-6, TLR4, and fecal LCN-2. These results highlight the critical role of microbial activity in modulating host responses to dietary fiber and suggest that proper fermentation of fibers is necessary to prevent accumulation of unfermented substrates that may provoke colonic inflammation.

**Fig 4.**
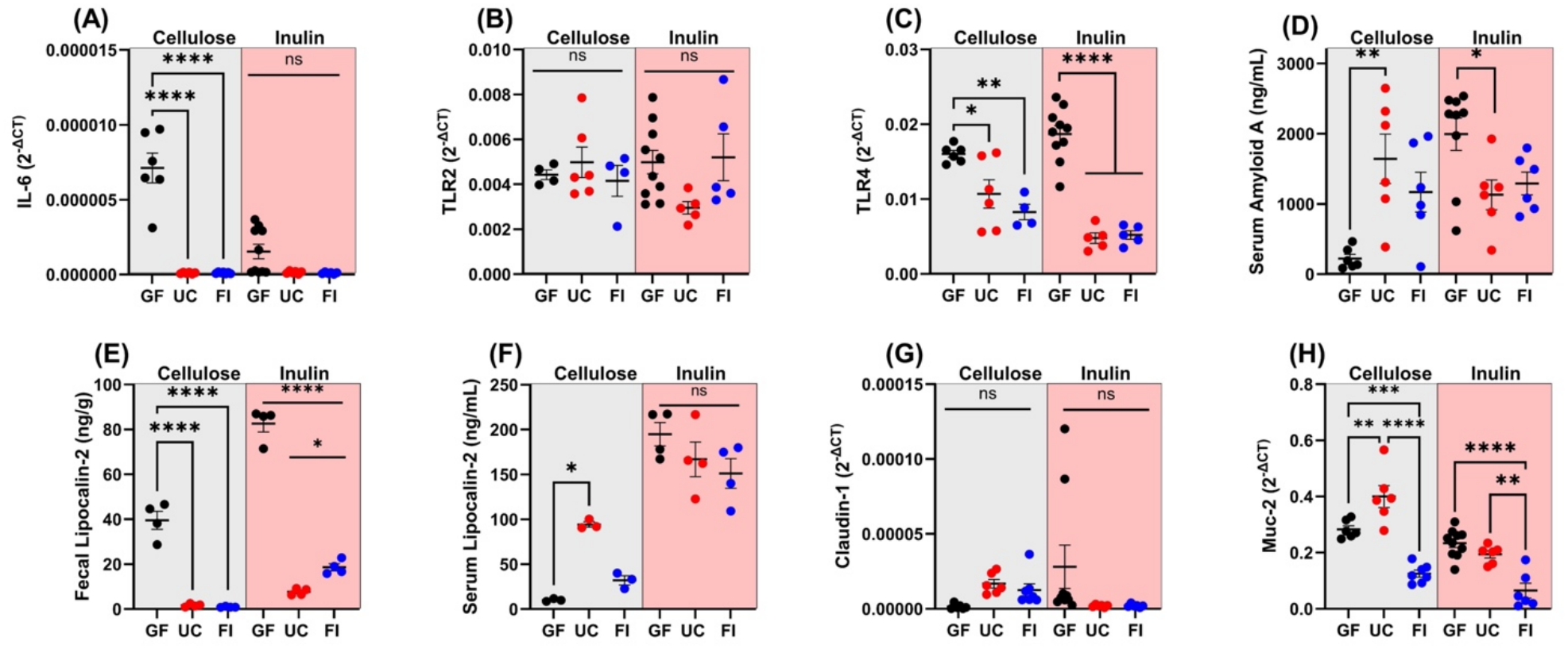
Relative gene expression of IL-6 (A), TLR2 (B), TLR4 (C), Serum Amyloid A (D), Lipocalin-2 in fecal (E) and serum (F), Claudin-1 (G), and Muc-2 (H) in mice fed with cellulose- and inulin-containing diets in GF, UC and FI colonized conditions. Replicates (n = 6–10) for each mice group are plotted. An asterisk (*) and horizontal line represent a statistical difference (p ≤ 0.05) between the two groups. (**) represents p ≤ 0.01, (**) represents p ≤ 0.001, (**) represents p ≤ 0.0001 and (ns) means no statistical difference. Data is shown as mean ± SEM. Alt text: Multi-panel figure showing relative gene expression of IL-6 (A), TLR2 (B), TLR4 (C), serum amyloid A concentrations (D), fecal (E) and serum (F) lipocalin-2 concentrations, claudin-1 (G) and Muc-2 (H) relative gene expression in GF mice and mice colonized with microbiota from UC and FI donors, fed cellulose-or inulin-containing diets.

### 3.3 Il10 deficiency is not the sole driving factor behind inulin-associated inflammation

To evaluate whether host genetic background contributes to inulin sensitivity, particularly in the context of a predisposition to spontaneous colitis, we compared GF Il10^⁻/⁻^ mice to GF WT mice with diets enriched with either cellulose or inulin. As shown in Figure 5, cellulose supplementation did not induce weight loss in either genotype (Fig. 5A). Notably, the WT mice exhibited a significant increase in body weight by the second week. In contrast, inulin diet fed group revealed a distinct genotype-dependent response (Fig. 5B); while WT mice maintained stable body weight throughout the study, Il10^⁻/⁻^ mice displayed progressive weight loss at both one and two weeks. These results suggest that inulin sensitivity, as measured by weight loss, is more pronounced in mice genetically predisposed to inflammatory bowel disease.

**Fig 5.**
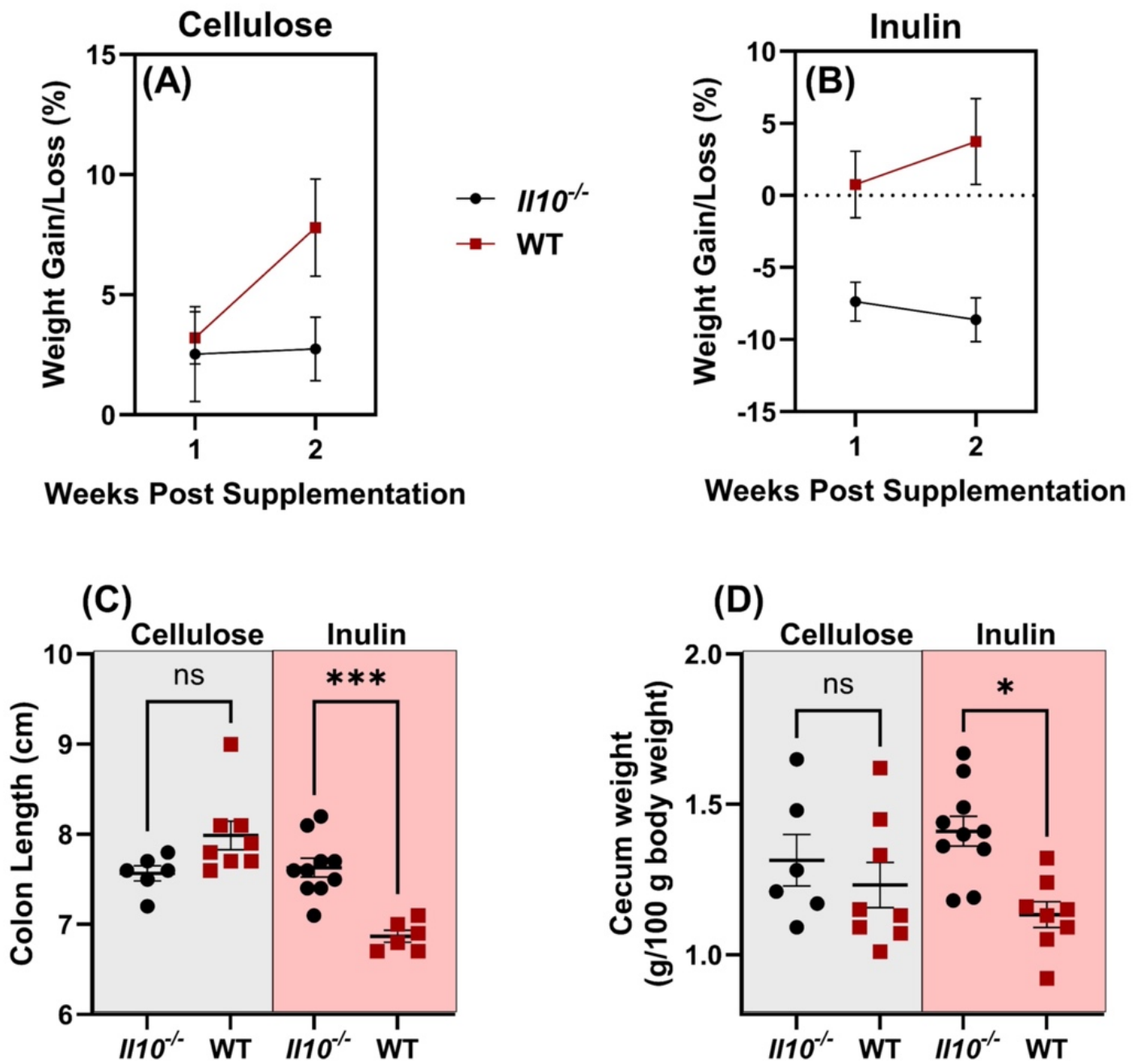
Body weight gain or loss percentage after one- or two-week supplementation with cellulose (A) or inulin (B) in Il10^⁻/⁻^ genetically modified and WT mice. Mouse colon lengths (C) and cecum weights (D) for cellulose and inulin fed groups in Il10⁻/⁻ genetically modified and WT mice. Replicates (n = 6–10) for each group are plotted. An asterisk (*) and horizontal line represent a statistical difference (p ≤ 0.05) between the two groups. (***) represents p ≤ 0.001 and (ns) means no statistical difference. Data is shown as mean ± SEM. Alt text: Multi-panel figure showing body weight gain or loss percentage (A-B), colon lengths (C), and cecum weights (D) in Il10⁻/⁻ genetically modified and WT mice fed cellulose- or inulin-containing diets.

Assessment of organ weights revealed genotype-dependent differences in colon length. In cellulose diet fed group, colon lengths were not significantly different between genotypes (Fig. 5C), whereas inulin fed Il10^⁻/⁻^ mice displayed increased colon length relative to the WT controls (Fig. 5C). This increase may reflect altered luminal accumulation and reduced fermentation of inulin in Il10^⁻/⁻^ mice, leading to cecal and colonic distension, potentially exacerbated by impaired immune regulation in this genotype. No significant differences were observed in colon weights, colon weight-to-length ratios, or in liver, spleen, and kidney weights across genotypes or fiber types (Fig. S4). Cecum weights were not different between genotypes under cellulose feeding; however, Il10^⁻/⁻^ mice exhibited significantly increased cecum weights following inulin supplementation (Fig. 5D).

Evaluation of inflammatory markers between genotypes revealed no significant differences in IL-6 expression under cellulose diet. However, inulin-fed WT mice exhibited higher colonic IL-6 expression compared to Il10^⁻/⁻^ mice (Fig. 6A). Expression of TLR2 (Fig. 6B) and TLR4 (Fig. 6C) did not differ significantly between genotypes under either dietary fiber groups. Additionally, WT mice showed higher expression of Claudin-1 (Fig. 6D) and Muc-2 (Fig. 6E) under both cellulose and inulin diets, suggesting enhanced maintenance of gut barrier integrity relative to the Il10^⁻/⁻^ counterparts.

**Fig 6.**
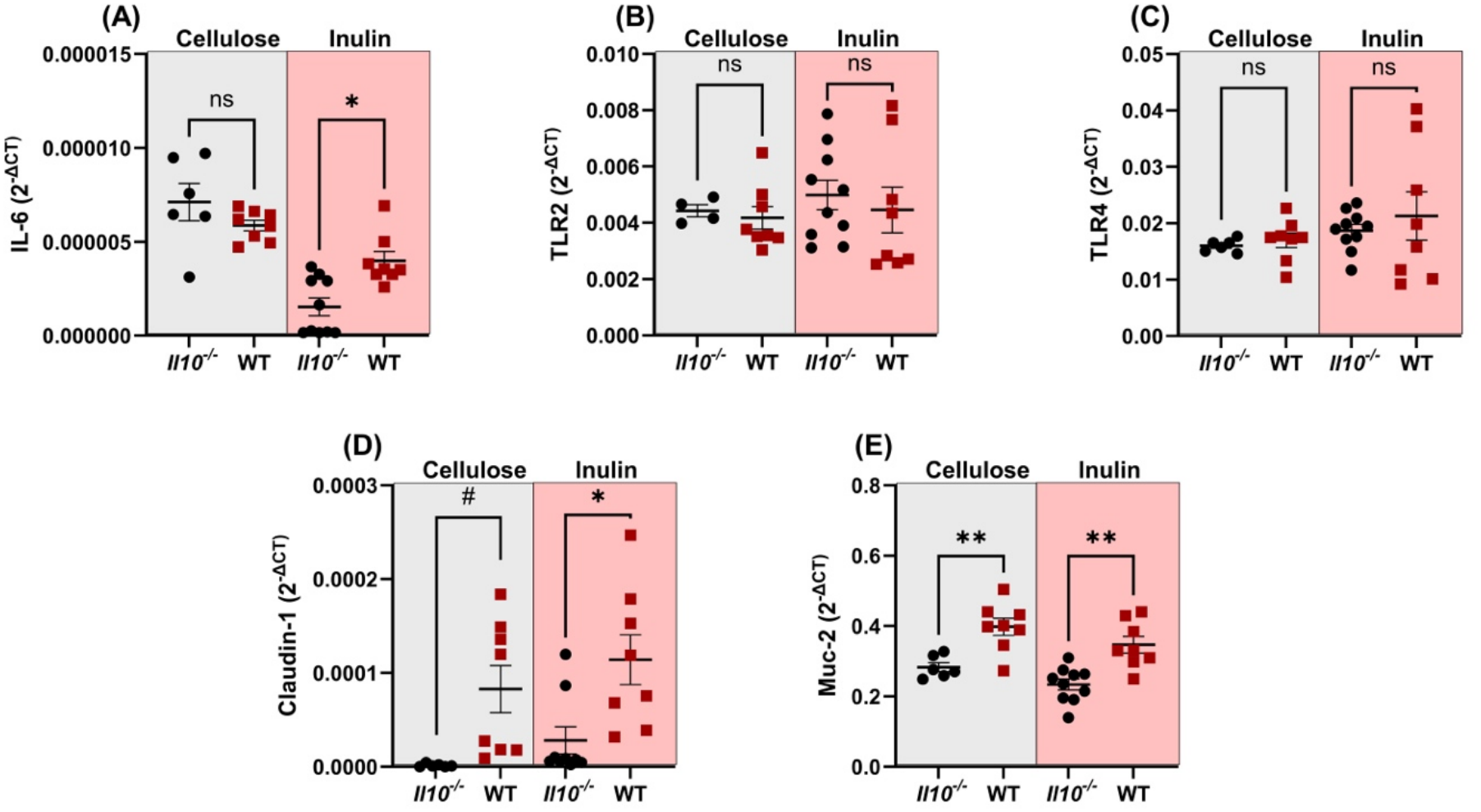
Relative gene expression of IL-6 (A), TLR2 (B), TLR4 (C), Claudin-1 (D), and Muc-2 (E) in distal colon tissues of mice fed with cellulose and inulin in Il10⁻/⁻ genetically modified and WT mice. Replicates (n = 6–10) for each mouse group are plotted. An asterisk (*) and horizontal line represent a statistical difference (p ≤ 0.05) between the two groups. (**) represent p ≤ 0.01, (#) indicates a trend toward statistical significance (0.05 < p ≤ 0.10), and (ns) means no statistical difference. Data is shown as mean ± SEM. Alt text: Multi-panel figure showing relative gene expression of IL-6 (A), TLR2 (B), TLR4 (C), claudin-1 (D), and Muc-2 (E) in distal colon tissues of Il10⁻/⁻ genetically modified and WT mice fed cellulose- or inulin-containing diets.

Overall, while dietary inulin-induced weight loss is more pronounced in Il10^⁻/⁻^ mice, other inflammatory and physiological responses to fiber are not consistently driven by host genotype alone. Despite their genetic predisposition to colitis, Il10^⁻/⁻^ mice did not exhibit elevated expression of pro-inflammatory markers compared to WT controls under most conditions. However, WT mice consistently displayed higher expression of gut barrier integrity markers, including Claudin-1 and Muc-2, across both fiber types. This may suggest that genetic predisposition is not the driving factor behind inulin intolerance , which underscores the the importance of the bacterial metabolism of fibers and their proper fermentation.

### 3.4 Fiber structure and chemical composition determine the inflammatory potential of inulin

To investigate whether chemical structure of dietary fibers contribute to host sensitivity, independently of genetic predisposition or microbial status, we compared the effects of several fibers with differing monomeric sugar residues but comparable chain length and size of the polymer. As shown in Figure 7A, fiber-specific differences in host response were observed, with fructooligosaccharides (FOS) inducing the most severe response. FOS resulted in mice with marked body weight loss by day 6, necessitating early euthanasia. Analysis of colonic inflammatory markers revealed that FOS and inulin from chicory (ChIn) supplementation significantly increased the expression of IL-6 as compared to the inulin from agave (AgIn)- and modified citrus pectin (MCP)-fed groups. Additionally, FOS supplementation resulted in increased expression of TLR2, TLR4, and TLR5 compared to other fiber treatments (Fig. 7C–E), indicating enhanced activation of host innate immune reponses. This response was accompanied by a modest increase in Claudin-1 (Fig. 7G). In contrast, dietary MCP did not elicit any pro-inflammatory response and significantly increased expression of Muc-2 (Fig. 7F), suggesting a protective role in maintaining epithelial barrier integrity. It is evident that FOS, a short-chain β-fructan, induced significant adverse effects, including severe weight loss and heightened expression of inflammatory markers, confirming its detrimental role in this model. Similarly, ChIn, a pool of slightly longer-chain β-fructans, also elicited elevated inflammatory responses, suggesting that fructan-type fibers may exacerbate inflammation under certain host conditions. In support of this, AgIn did not induce inflammatory responses, highlighting that subtle structural differences among fructan-type fibers (here, branched vs. linear) can lead to distinct host physiological and immune response outcomes. Some of these effects may be partly attributable to the presence of difructose anhydride III (DFAIII) in ChIn, a byproduct of its processing that has been reported to resist microbial fermentation and inhibit bacterial bile acid 7α-dehydroxylase activity, potentially amplifying immune activation in the colonic environment^30,31^. In contrast, agave-derived inulin, which consists predominantly of linear FF (fructose-fructose)-type fructan chains lacking a terminal glucose residue and DFAIII-type anhydrides, did not elicit pro-inflammatory responses, suggesting that these structural and composition distinctions, rather than fructan chain length or linkage patterns alone, may govern the immunological outcome of dietary fiber supplementation in susceptible hosts.

**Fig 7.**
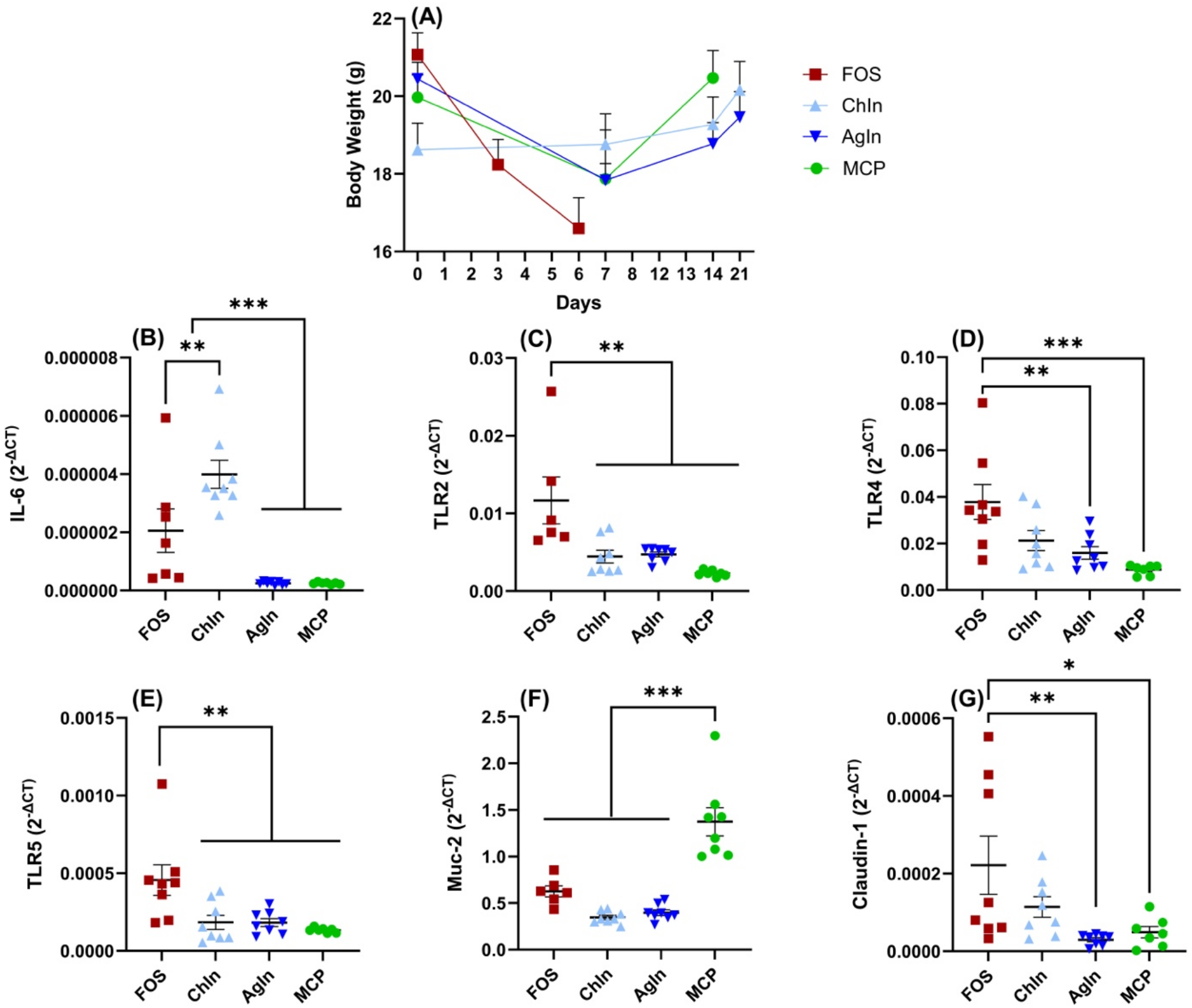
Body weights throughout the experimental period of mice fed with fructooligosaccharides (FOS), ChIn (inulin from chicory), AgIn (inulin from agave), and modified citrus pectin (MCP) (A). Relative gene expression of IL-6 (B), TLR2 (C), TLR4 (D), TLR5 (E), Muc-2 (F), and Claudin-1 (G) in distal colon tissues of GF WT mouse fed with distinct high-fiber diets. Replicates (n = 6–10) for each fiber treatment are plotted. An asterisk (*) and horizontal line represent a statistical difference (p ≤ 0.05) between the two groups. (**) represents p ≤ 0.01, and (**) represents p ≤ 0.001. Data is shown as mean ± SEM. Alt text: Multi-panel figure showing body weights (A) and relative gene expression of IL-6 (B), TLR2 (C), TLR4 (D), TLR5 (E), Muc-2 (F), and claudin-1 (G) in distal colon tissues of GF WT mice fed with fructooligosaccharides (FOS), ChIn (inulin from chicory), AgIn (inulin from agave), or modified citrus pectin (MCP).

## 4. Discussion

Dietary fiber is widely recognized for its role in promoting gut health, supporting gut microbial diversity, and reducing systemic inflammation^32^. Yet paradoxically, patients with IBD frequently report adverse gastrointestinal responses to fiber supplementation, despite these well-established benefits^33^. In this study, we demonstrate that the absence of microbial fermentation is a critical determinant of fiber intolerance, as GF Il10⁻/⁻ mice developed severe inflammatory responses specifically in response to inulin supplementation, extending prior *in vitro* and *ex vivo* observations showing that unfermented β-fructans can provoke inflammation in susceptible individuals.

In GF Il10⁻/⁻ mice, inulin induced pronounced weight loss, diarrhea, rectal prolapse, and morphological alterations including cecal distension and colonic thinning, whereas these effects were entirely absent in CONV Il10⁻/⁻ mice, underscoring the protective role of an intact and functional microbiome. The accumulation of intact inulin in the cecum, confirmed by elevated fructose concentrations following acid hydrolysis, supports the hypothesis that unfermented fiber exerts osmotic and mechanical stress on the intestinal epithelium (Fig. 1), consistent with observations in GF models where unfermented fiber polymers accumulate and disrupt luminal homeostasis. Beyond osmotic effects, unfermented β-fructans may directly engage epithelial and mucosal immune cells independently of microbial metabolism.

Histological examination of Alcian blue-stained distal colon sections provided compelling morphological evidence of this mucosal compromise. Cellulose-fed mice maintained a robust, continuous mucus layer with abundant goblet cell content, whereas inulin-fed GF mice exhibited marked depletion of Alcian blue staining suggesting reduced goblet cell population or Muc2 expression (Fig. 1J–K), indicating disruption of the mucus barrier. This structural deterioration likely increases epithelial exposure to luminal contents, creating conditions permissive for immune activation and reinforcing the molecular findings described below.

At the molecular level, upregulation of TLR2 and TLR4 in GF inulin-fed mice points to direct activation of host innate immune receptors (Fig. 2). While TLRs are classically associated with pattern recognition receptors (PRR), accumulating evidence indicates that β-fructans themselves can modulate TLR signaling in a structure-dependent manner. β (2→1)-fructans have been shown to activate TLR2 and influence epithelial barrier function in a chain length-dependent fashion, whereas fructans can either promote or attenuate inflammation depending on their structural configuration and degree of polymerization. In the absence of gut microbiota, gut epithelial stress and damage-associated molecular patterns (DAMPs), including ATP release, heat shock proteins, and host-derived extracellular matrix fragments generated under osmotic or mechanical stress, may serve as endogenous TLR ligands, shifting the balance toward a pro-inflammatory state. This is consistent with previous work^34^, which established that TLR signaling is not merely a response to microbial products but is fundamentally required for maintaining gut homeostasis, and that dysregulation of this innate immune signaling, whether by microbe-associated or damage-associated signals, promotes intestinal inflammatory disease. Elevated circulating SAA and LCN-2 in GF inulin-fed mice further confirm that local intestinal inflammation extends systemically (Fig. 2) potentially via ‘leaky gut’, suggesting that unfermented inulin not only disrupts epithelial homeostasis but propagates systemic inflammatory responses in genetically susceptible hosts.

To assess whether microbial communities could modulate these effects, we engrafted microbiota from UC and FI donors into GF Il10⁻/⁻ mice. Even dysbiotic microbe colonization partially mitigated inulin-induced weight loss and reduced key inflammatory markers (Fig. 3–4), demonstrating that microbial metabolic activity is the primary protective factor. Notably, even the minimal microbial presence conferred by colonization with these dysbiotic/altered communities was sufficient to partially restore fiber tolerance, reminiscent of findings showing that defined minimal consortia such as the Altered Schaedler Flora (ASF) can be sufficient to prime appropriate immune responses and prevent exaggerated innate immune activation in germ-free settings^35^. This aligns with Rakoff-Nahoum *et al.* work^34^ that commensal-derived PRR signals are required for proper gut epithelial and immune education, and that their complete absence predisposes to dysregulated responses. The incomplete recovery in colonized mice, however, suggests that not all microbiota confer equal protection, dysbiotic communities may lack key fiber-degrading taxa or produce altered metabolic outputs leading to inefficient fermentation. Reduced TLR4 expression and fecal LCN-2 in colonized mice, even from diseased donors, suggest that any degree of microbial presence attenuates host innate immune overactivation (Fig. 4). The differential modulation of Muc-2 by colonization source further reinforces this point: although UC-derived microbiota enhanced Muc-2 expression, FI-derived microbiota consistently suppressed it across fiber types (Fig. 4H), suggesting that certain dysfunctional communities actively impair mucus production and compromise barrier integrity, potentially increasing epithelial permeability to luminal antigens. These findings are particularly relevant to human IBD, where reduced abundance of butyrate-producing taxa such as *Faecalibacterium prausnitzii* and *Roseburia* spp. has been linked to impaired barrier function and increased inflammation. Taken together, these results highlight that the functional capacity of the microbiota, rather than its mere presence, is the critical determinant of host responses to dietary fiber.

Regarding host genetics, whereas Il10⁻/⁻ mice exhibited greater inulin-induced weight loss than WT counterparts (Fig. 5), inflammatory markers were not consistently elevated between genotypes (Fig. 6). Interestingly, WT mice showed higher expression of barrier-associated genes including Claudin-1 and Muc-2, suggesting that intact immune regulation enhances epithelial resilience independent of inflammatory tone. The increased cecal weight and colon length in Il10⁻/⁻ mice likely reflect luminal distension from unfermented polymers, gas production by chemical fermentation accumulation rather than a genotype-driven inflammatory response per se. These findings indicate that genetic predisposition alone is insufficient to drive inulin-induced gut inflammation, rather, it operates as a sensitizing factor within a broader interplay of microbial metabolism and fiber structure.

Perhaps the most striking finding of this study is that inflammatory responses to inulin are not a class effect, but depend on fiber source, chemical structure, and composition. Among structurally matched fibers, FOS and ChIn drove the strongest inflammatory responses, AgIn and MCP were well tolerated (Fig. 7). This divergence is not arbitrary; it reflects fundamental chemical differences between these fibers that have profound immunological consequences.

Chicory inulin is a β (2→1)-linked fructan with a terminal glucose residue and, critically, contains DFAIII, a byproduct generated during thermal or enzymatic processing. DFAIII has been reported to resist microbial fermentation and inhibit bacterial bile acid 7α-dehydroxylase activity, an enzyme critical for secondary bile acid generation^30,31^. This inhibition may disrupt bile acid signaling pathways (FXR, TGR5) that normally exert anti-inflammatory effects in the colon, while simultaneously allowing intact fructan structures to persist and engage innate immune receptors.

FOS, as a short-chain β (2→1)-fructan, shares this structural backbone but with a lower degree of polymerization, which may enhance its direct accessibility to epithelial TLR binding sites and explain the most severe inflammatory phenotype observed in this model. AgIn, by contrast, presents a fundamentally different chemical structure. Its composed predominantly of branched FF-type fructan chains (fructose-fructose linkages without a terminal glucose residue), has a higher average degree of polymerization, and lacks the DFAIII-type anhydride byproducts characteristic of processed chicory inulin. Its slower, and more uniform fermentation profile and distinct linkage chemistry may limit direct engagement with PRRs, explaining the absence of inflammatory responses observed in our AgIn-fed mice. This distinction between two fibers both classified as “inulin” illustrates that fiber alone is an inadequate guide for clinical recommendations. MCP further reinforces this point from a different structural angle: its complex, branched polygalacturonic acid backbone has no structural resemblance to fructan-type receptor ligands, likely limiting direct TLR engagement while actively promoting Muc-2 expression and barrier support (Fig. 7F), consistent with prior work demonstrating pectin’s protective properties over inulin in gut inflammatory contexts^36^.

Some limitations of this study include the duration of microbial colonization, which was relatively short and may not fully reconstitute microbial ecological complexity or fermentative capacity. The use of a limited number of human donors also restricts generalizability. Future studies incorporating defined microbial consortia, metabolomic profiling, and extended colonization periods will be essential to further dissect the mechanisms underlying fiber-specific immune responses.

In conclusion, our findings establish microbial fermentation as the central gatekeeper of host tolerance to dietary fermentable fiber. In its absence, inulin acts as a direct pro-inflammatory stimulus, triggering mucus depletion, innate immune activation via TLR signaling, and systemic inflammation - effects that are partially rescued even by dysbiotic microbiota colonization, reinforcing that fermentative activity, however imperfect, is protective. Critically, fiber structure is not a secondary consideration but a primary determinant of immunological outcome: chicory and agave inulin, despite sharing a fructan identity, produce opposite responses driven by differences in linkage chemistry, terminal residues, and processing-derived anhydride content. These findings carry direct clinical implications; dietary fiber recommendations for individuals with IBD, fiber intolerance, or gut microbial dysbiosis must move beyond generic fiber categories and instead account for the structural identity of the fiber, the fermentative capacity of the resident microbiota, and the immune status of the host.

## Funding

The funders had no role in the study design, collection, or interpretation of data. This research is funded by the United States Department of Agriculture Foundation grant 2022-67017-37184 and Whistler Center for Carbohydrate Research and Heartland Foundation grants.

## Conflicts of Interest

The authors disclose no conflicts.

## Conference Presentation

Part of this work was presented at Digestive Disease Week 2026 and NUTRITION 2026.

## Acknowledgments

We thank the patients (donors) of the study for agreeing to participate and provide samples, as well as the research coordinators, nurses and staff at The Seattle’s Children Hospital. We acknowledge the Purdue Gnotobiotic Core and the Histology Research Laboratory at Purdue University.

## Ethical Considerations

Human fecal sample collection was approved by Purdue’s Institutional Review Board (IRB-2021-109), and consent was obtained from patients/guardian. All animal procedures were approved by the Purdue Institutional Animal Care and Use Committee (PIACUC approval No. 1810001817) according to the recommendations of the Guide for the Care and Use of Laboratory Animals published by the National Institutes of Health.

## Author Contributions

The authors confirm contributions to the paper as follows: E.M. study design, data acquisition, analysis and data interpretation, article drafting; P.A. data acquisition, analysis and data interpretation, article revision; V.N .data acquisition, analysis and data interpretation; Y.E.T data acquisition, analysis and data interpretation, article revision; S.R.L. analysis and data interpretation, and article revision; B.S.Y data acquisition, analysis and data interpretation, and article revision; A.T.G study design, article revision; T.W.L.C study design, analysis and data interpretation, article revision; M.V.K. study design, analysis and data interpretation, article revision; L.R. study design, data acquisition, analysis and data interpretation, article drafting and revision.

## Data Availability Statement

The data underlying this article are available in the article and in its online supplementary material.

---

**Fig S1.**
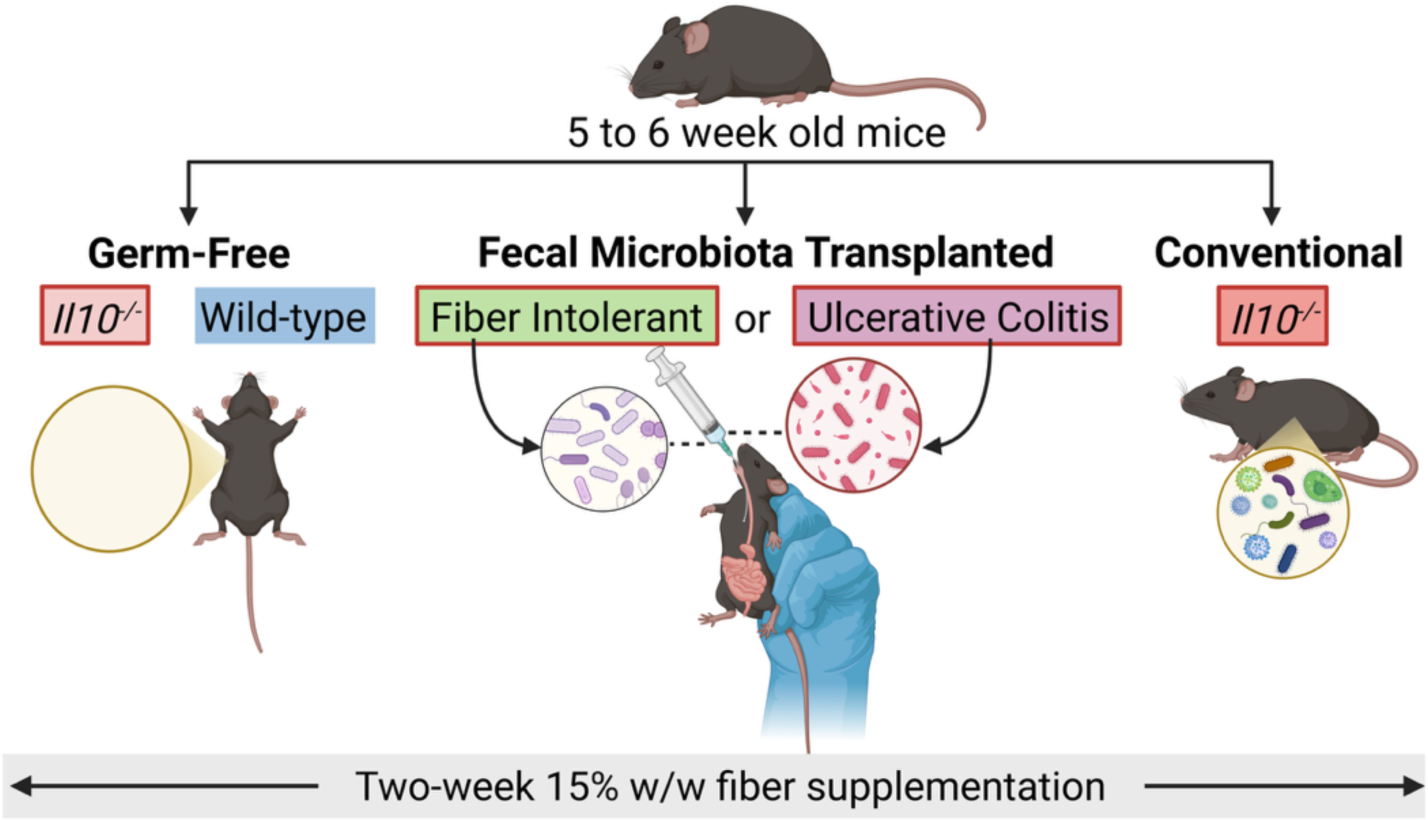
Experimental Design. Mice were randomly assigned to dietary treatment groups, with an equal number of males and females allocated to each experimental condition. Fecal microbiota transplanted mice with UC and FI microbiota were Il10^-/-^.

**Fig S2.**
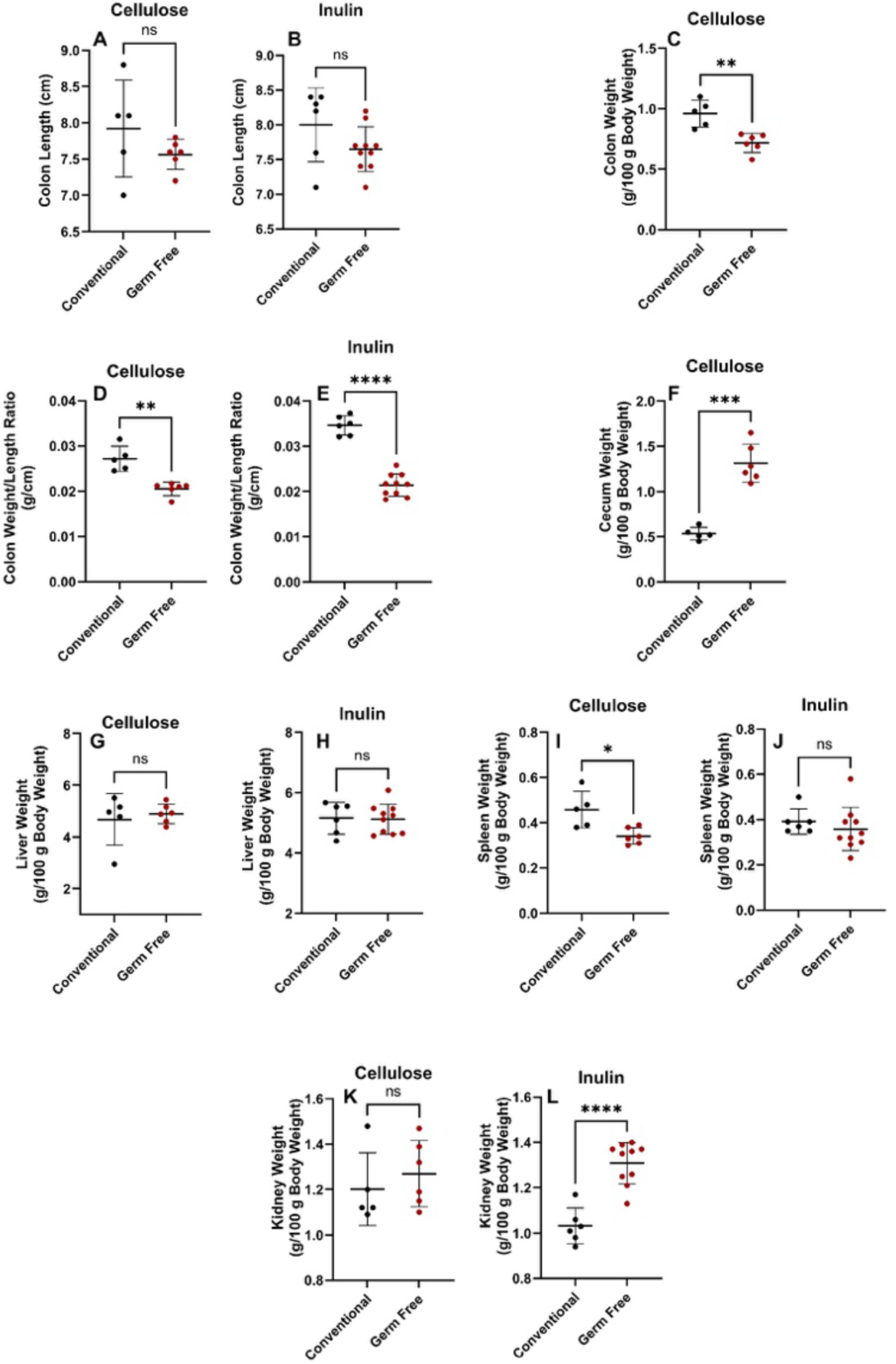
Mice organ weights and colon lengths for cellulose and inulin treatments in conventional and germ-free conditions. Replicates (n = 6–10) for each mice group are plotted. An asterisk (*) and horizontal line represent a statistical difference (p ≤ 0.05) between the two groups. (**) represents p ≤ 0.01, (**) represents p ≤ 0.001, (**) represents p ≤ 0.0001 and (ns) means no statistical difference. Data is shown as mean ± SEM.

**Fig S3.**
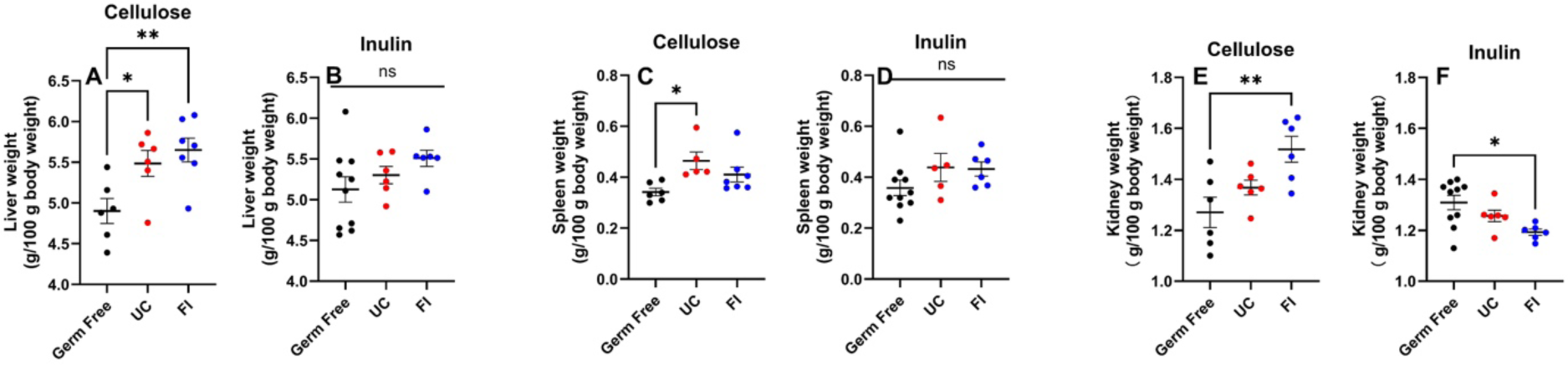
Mice organ weights and colon lengths for cellulose and inulin treatments in germ-free, and UC and FI colonized conditions. Replicates (n = 6–10) for each mice group are plotted. An asterisk (*) and horizontal line represent a statistical difference (p ≤ 0.05) between the two groups. (**) represents p ≤ 0.01, (**) represents p ≤ 0.001, (**) represents p ≤ 0.0001 and (ns) means no statistical difference. Data is shown as mean ± SEM.

**Fig S4.**
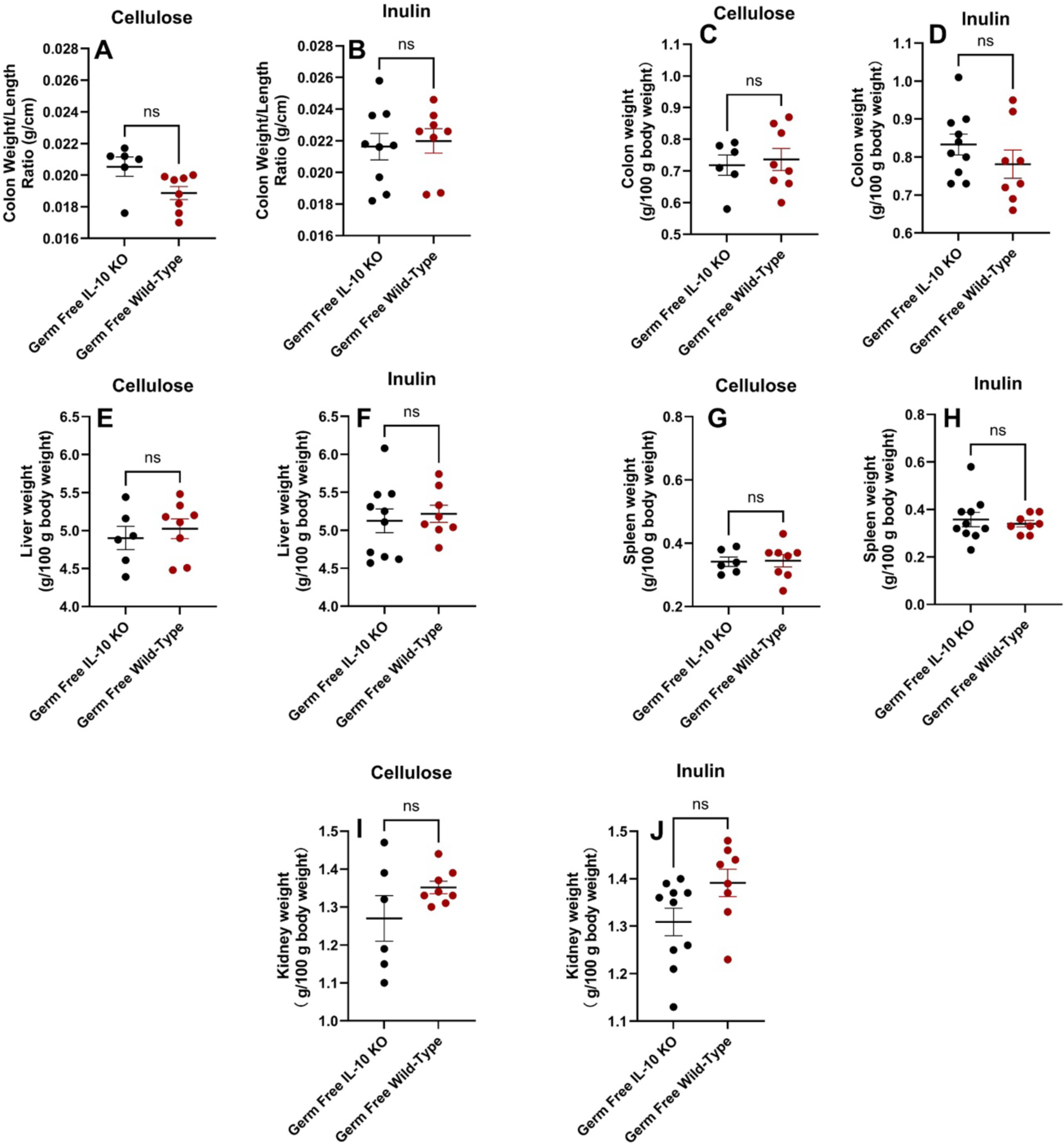
Mice organ weights and colon lengths for cellulose and inulin treatments in Il10⁻/⁻ genetically modified and wild-type C57BL/6J mice. Replicates (n = 6–10) for each mice group are plotted. An asterisk (*) and horizontal line represent a statistical difference (p ≤ 0.05) between the two groups. (***) represents p ≤ 0.001 and (ns) means no statistical difference. Data is shown as mean ± SEM.

**Fig S5.**
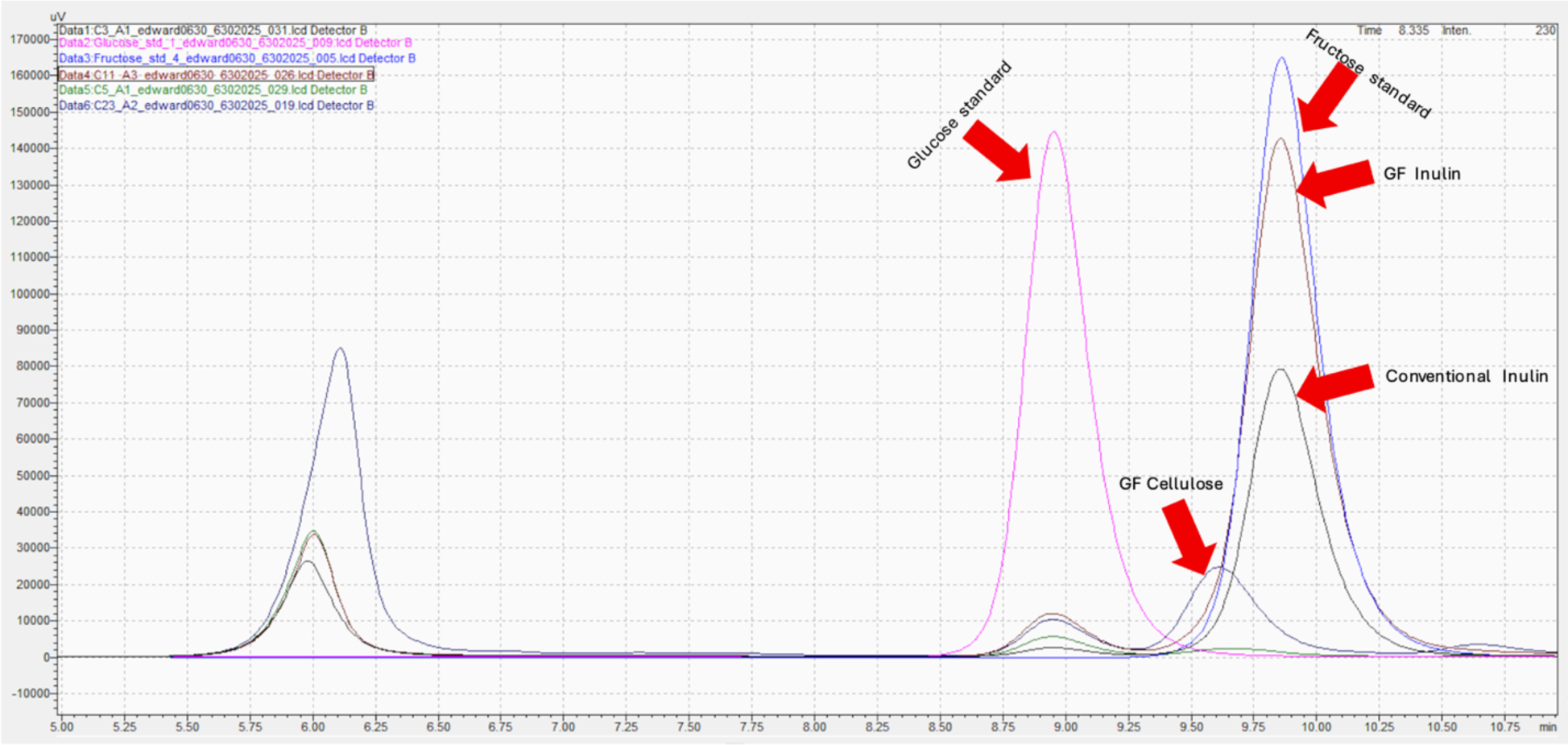
Representative HPLC chromatogram showing separation and detection of simple sugars in standards and experimental samples. The chromatogram includes glucose and fructose standards, as well as cecal digesta samples from germ-free (GF) and conventional (Conv) mice fed inulin- or cellulose-containing diets. Distinct fructose peaks are observed in the fructose standard and in samples from inulin-fed groups (GF and Conv), consistent with the presence of hydrolyzed inulin-derived fructose. No detectable fructose peak was observed in samples from Conv cellulose-fed groups, confirming the absence of fructose in these controls. Retention times of standards were used to identify corresponding peaks in experimental samples.

**Table S1.**
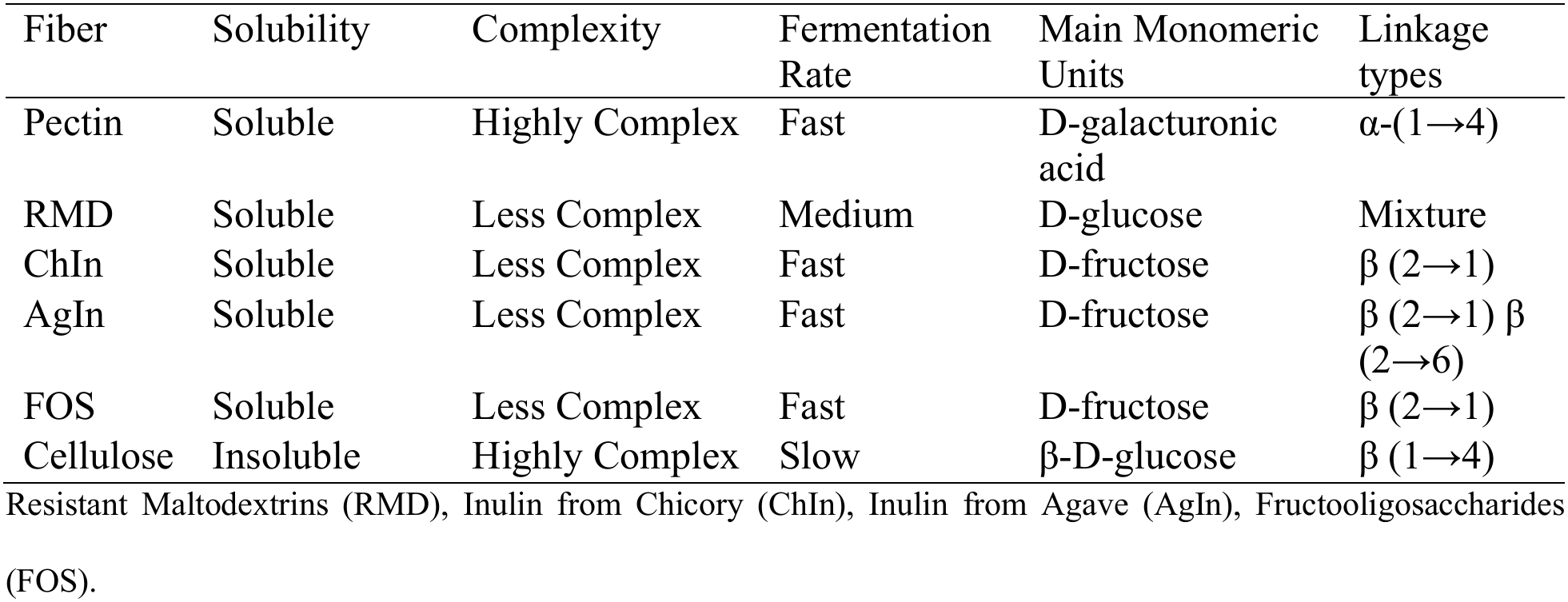
Physicochemical properties of the dietary fibers selected.

**Table S2.**
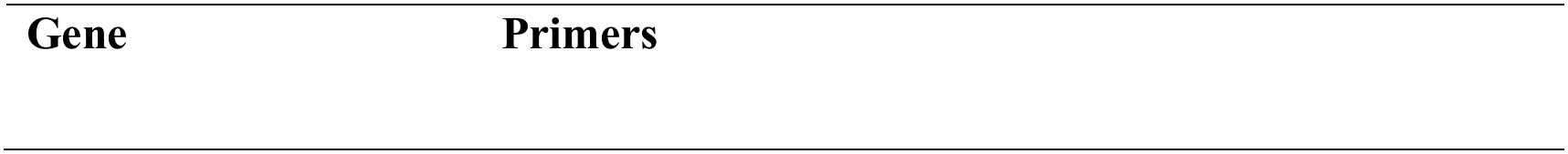
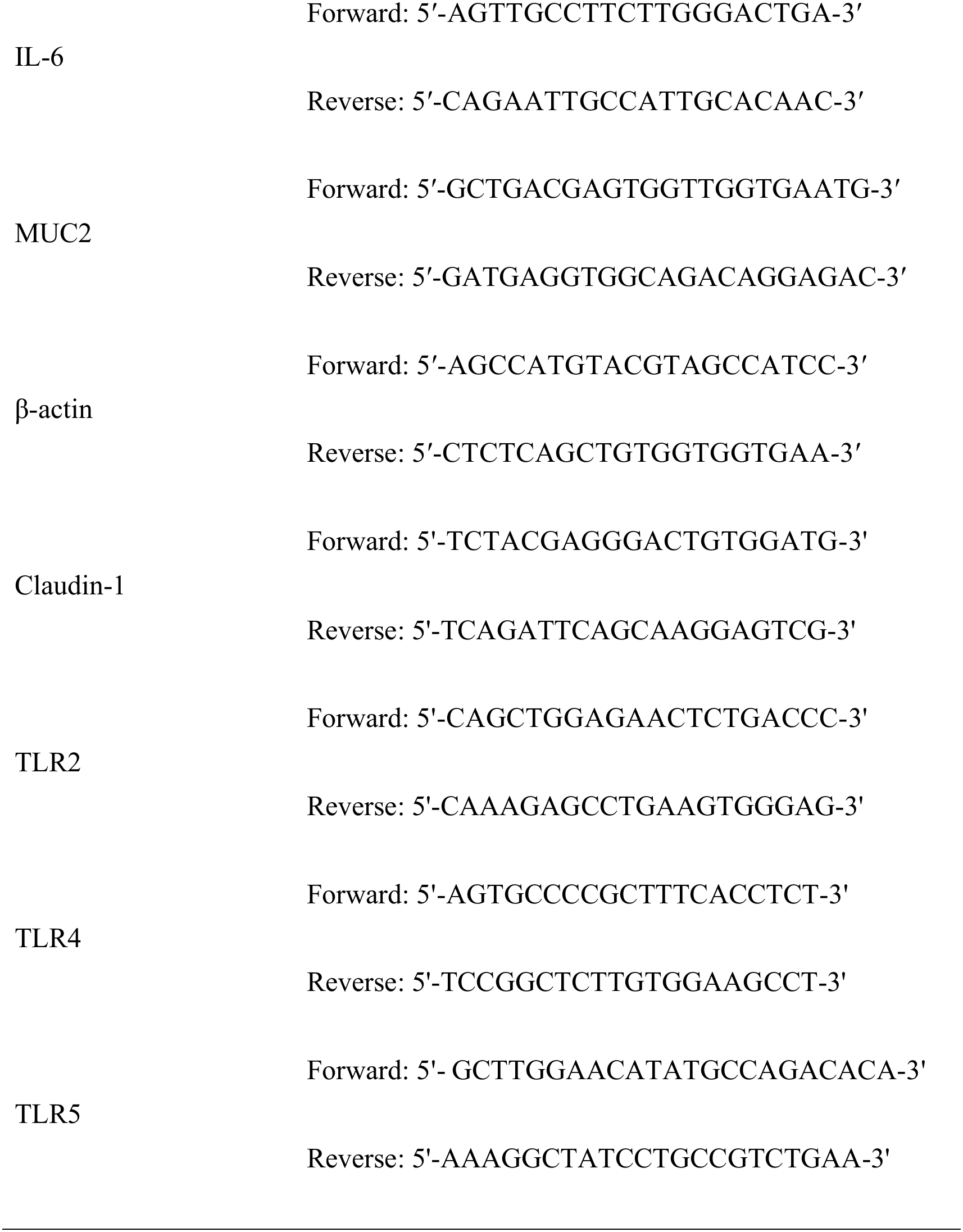
Real-time PCR primers.

**Table S3.**
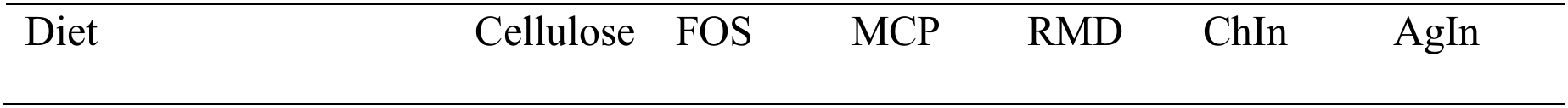
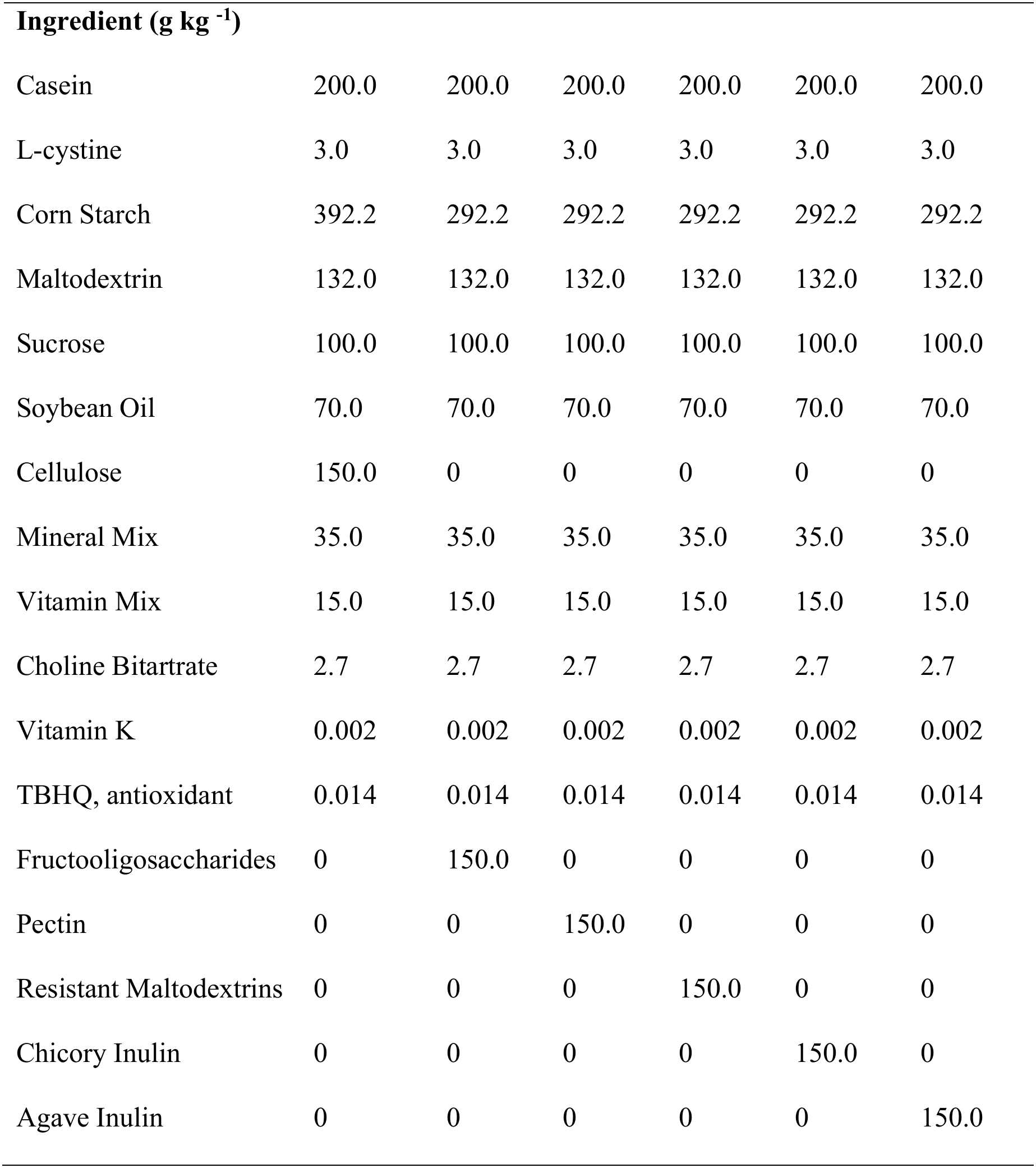
Composition of animal experiment diets.

## Notes

### Competing Interest Statement

The authors have declared no competing interest.

