## Supplementary Data for "Impaired Fermentation in the Absence of Gut Microbiota Promotes Inulin-Induced Intestinal Inflammation"

**Running Title:** Impaired Inulin Fermentation Promotes Gut Inflammation

***Corresponding Author:**

Lavanya Reddivari, PhD

Department of Food Science, Purdue University

745 Agriculture Mall Drive

West Lafayette, IN 47906, USA

**Funding:**

The funders had no role in the study design, collection, or interpretation of data. This research is funded by the United States Department of Agriculture Foundation grant 2022-67017-37184 and Whistler Center for Carbohydrate Research and Heartland Foundation grants.

**Conflicts of Interest:**

The authors disclose no conflicts.

**Summary:**

This study shows that microbial fermentation is essential for host tolerance to dietary fibers, and that fiber structure determines the immunological response.

**Supplementary Data**

**Table S1.** Physicochemical properties of the dietary fibers selected.

| Fiber | Solubility | Complexity | Fermentation Rate | Main Monomeric Units | Linkage types |
| --- | --- | --- | --- | --- | --- |
| Pectin | Soluble | Highly Complex | Fast | D-galacturonic acid | α-(1→4) |
| RMD | Soluble | Less Complex | Medium | D-glucose | Mixture |
| ChIn | Soluble | Less Complex | Fast | D-fructose | β (2→1) |
| AgIn | Soluble | Less Complex | Fast | D-fructose | β (2→1) β (2→6) |
| FOS | Soluble | Less Complex | Fast | D-fructose | β (2→1) |
| Cellulose | Insoluble | Highly Complex | Slow | β-D-glucose | β (1→4) |

Resistant Maltodextrins (RMD), Inulin from Chicory (ChIn), Inulin from Agave (AgIn), Fructooligosaccharides (FOS).

**Table S2.** Real-time PCR primers.

| **Gene** | **Primers** |
| --- | --- |
| IL-6 | Forward: 5′-AGTTGCCTTCTTGGGACTGA-3′ |
|  | Reverse: 5′-CAGAATTGCCATTGCACAAC-3′ |
| MUC2 | Forward: 5′-GCTGACGAGTGGTTGGTGAATG-3′ |
|  | Reverse: 5′-GATGAGGTGGCAGACAGGAGAC-3′ |
| β-actin | Forward: 5′-AGCCATGTACGTAGCCATCC-3′ |
|  | Reverse: 5′-CTCTCAGCTGTGGTGGTGAA-3′ |
| Claudin-1 | Forward: 5'-TCTACGAGGGACTGTGGATG-3' |
|  | Reverse: 5'-TCAGATTCAGCAAGGAGTCG-3' |
| TLR2 | Forward: 5'-CAGCTGGAGAACTCTGACCC-3' |
|  | Reverse: 5'-CAAAGAGCCTGAAGTGGGAG-3' |
| TLR4 | Forward: 5'-AGTGCCCCGCTTTCACCTCT-3' |
|  | Reverse: 5'-TCCGGCTCTTGTGGAAGCCT-3' |
| TLR5 | Forward: 5'- GCTTGGAACATATGCCAGACACA-3' |
|  | Reverse: 5'-AAAGGCTATCCTGCCGTCTGAA-3' |

**Table S3**. Composition of animal experiment diets.

| Diet | Cellulose | FOS | MCP | RMD | ChIn | AgIn |
| --- | --- | --- | --- | --- | --- | --- |
| **Ingredient (g kg ^-1^)** |  |  |  |  |  |  |
| Casein | 200.0 | 200.0 | 200.0 | 200.0 | 200.0 | 200.0 |
| L-cystine | 3.0 | 3.0 | 3.0 | 3.0 | 3.0 | 3.0 |
| Corn Starch | 392.2 | 292.2 | 292.2 | 292.2 | 292.2 | 292.2 |
| Maltodextrin | 132.0 | 132.0 | 132.0 | 132.0 | 132.0 | 132.0 |
| Sucrose | 100.0 | 100.0 | 100.0 | 100.0 | 100.0 | 100.0 |
| Soybean Oil | 70.0 | 70.0 | 70.0 | 70.0 | 70.0 | 70.0 |
| Cellulose | 150.0 | 0 | 0 | 0 | 0 | 0 |
| Mineral Mix | 35.0 | 35.0 | 35.0 | 35.0 | 35.0 | 35.0 |
| Vitamin Mix | 15.0 | 15.0 | 15.0 | 15.0 | 15.0 | 15.0 |
| Choline Bitartrate | 2.7 | 2.7 | 2.7 | 2.7 | 2.7 | 2.7 |
| Vitamin K | 0.002 | 0.002 | 0.002 | 0.002 | 0.002 | 0.002 |
| TBHQ, antioxidant | 0.014 | 0.014 | 0.014 | 0.014 | 0.014 | 0.014 |
| Fructooligosaccharides | 0 | 150.0 | 0 | 0 | 0 | 0 |
| Pectin | 0 | 0 | 150.0 | 0 | 0 | 0 |
| Resistant Maltodextrins | 0 | 0 | 0 | 150.0 | 0 | 0 |
| Chicory Inulin | 0 | 0 | 0 | 0 | 150.0 | 0 |
| Agave Inulin | 0 | 0 | 0 | 0 | 0 | 150.0 |

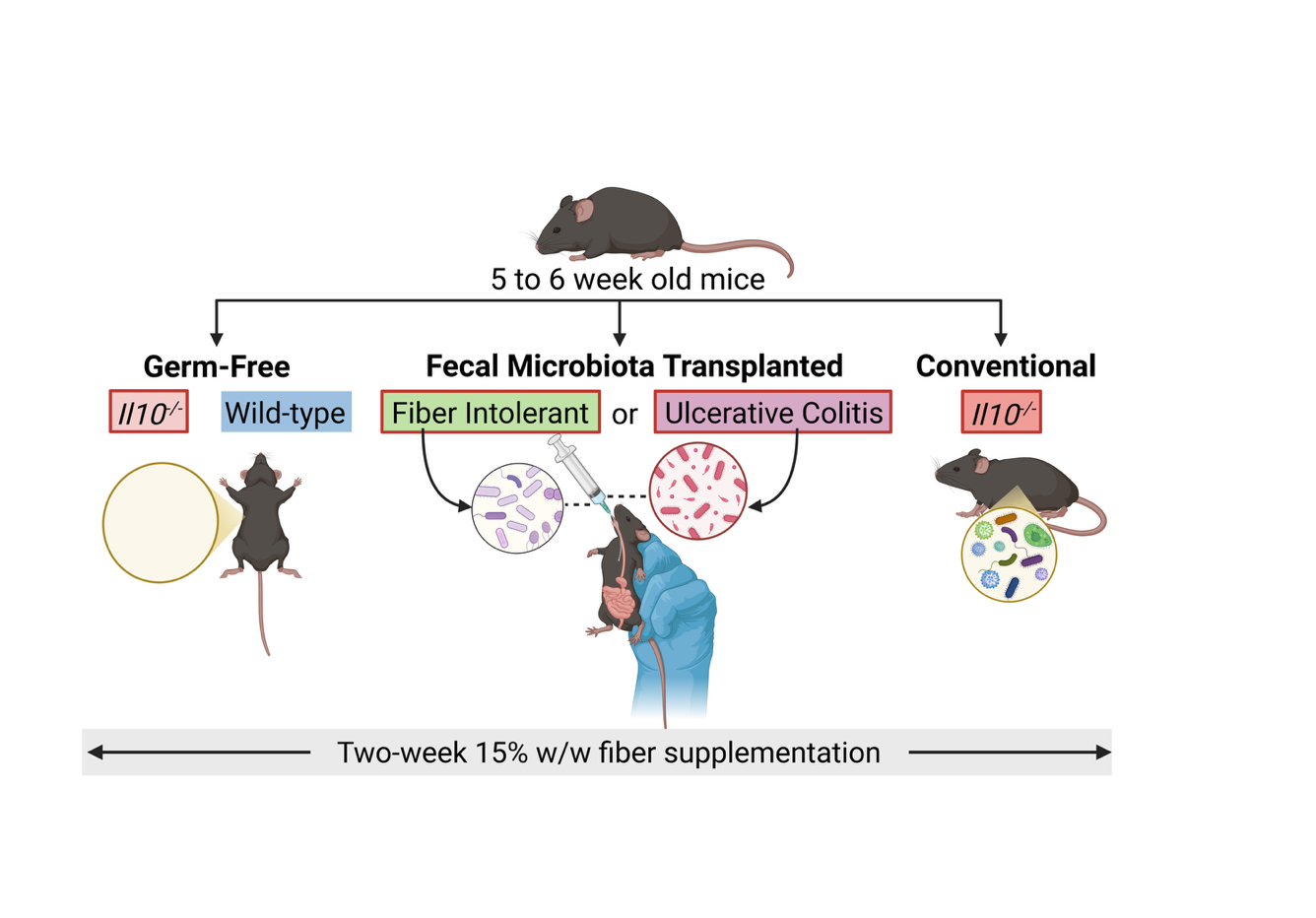

**Fig S1.** Experimental Design. Mice were randomly assigned to dietary treatment groups, with an equal number of males and females allocated to each experimental condition. Fecal microbiota transplanted mice with UC and FI microbiota were Il10^-/-^.

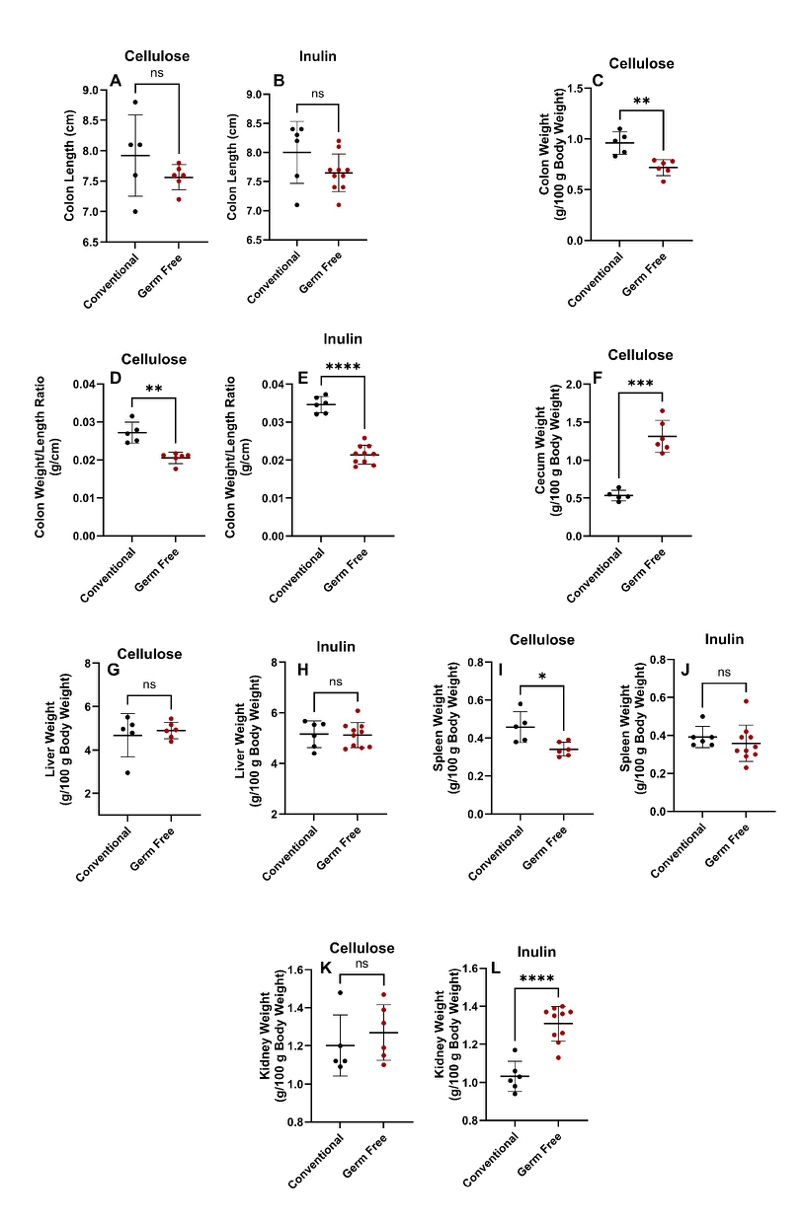

**Fig S2**. Mice organ weights and colon lengths for cellulose and inulin treatments in conventional and germ-free conditions. Replicates (n = 6–10) for each mice group are plotted. An asterisk (*) and horizontal line represent a statistical difference (p ≤ 0.05) between the two groups. (**) represents p ≤ 0.01, (**) represents p ≤ 0.001, (**) represents p ≤ 0.0001 and (ns) means no statistical difference. Data is shown as mean ± SEM.

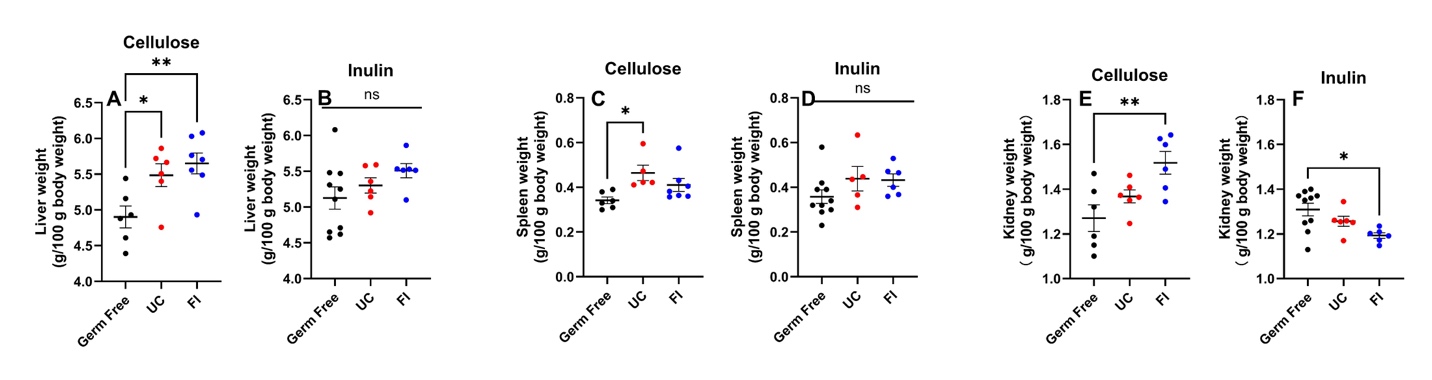

**Fig S3**. Mice organ weights and colon lengths for cellulose and inulin treatments in germ-free, and UC and FI colonized conditions. Replicates (n = 6–10) for each mice group are plotted. An asterisk (*) and horizontal line represent a statistical difference (p ≤ 0.05) between the two groups. (**) represents p ≤ 0.01, (**) represents p ≤ 0.001, (**) represents p ≤ 0.0001 and (ns) means no statistical difference. Data is shown as mean ± SEM.

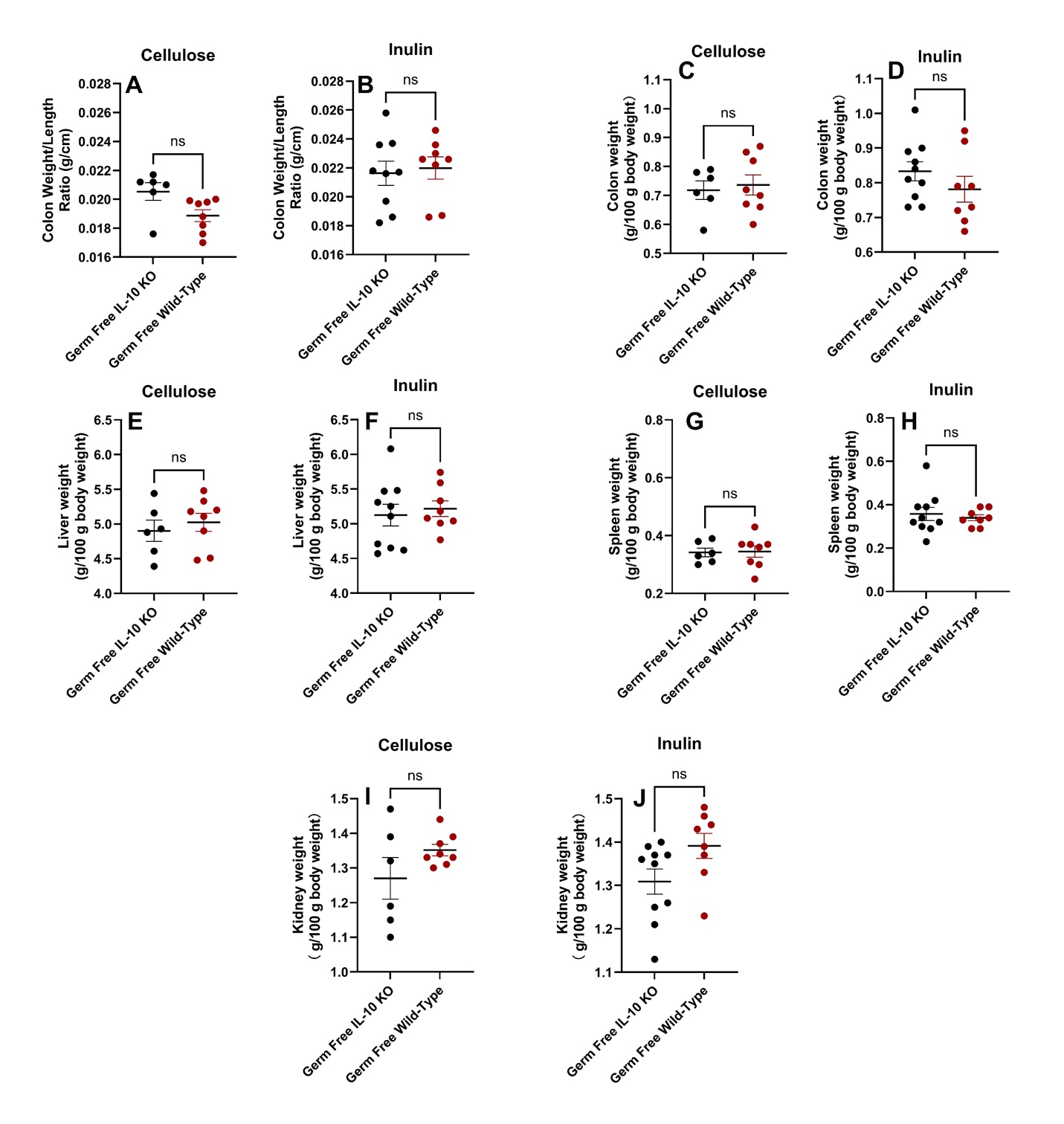

**Fig S4**. Mice organ weights and colon lengths for cellulose and inulin treatments in Il10⁻/⁻ genetically modified and wild-type C57BL/6J mice. Replicates (n = 6–10) for each mice group are plotted. An asterisk (*) and horizontal line represent a statistical difference (p ≤ 0.05) between the two groups. (***) represents p ≤ 0.001 and (ns) means no statistical difference. Data is shown as mean ± SEM.

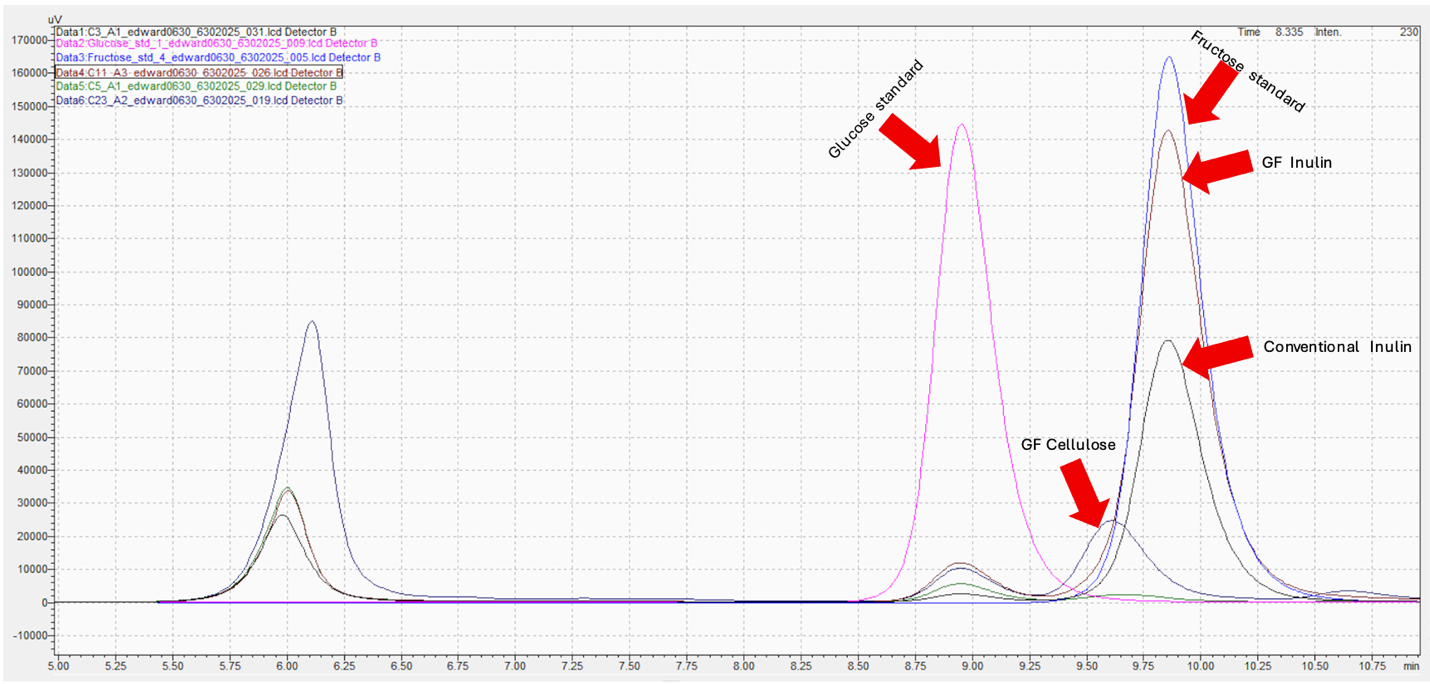

**Fig S5**. Representative HPLC chromatogram showing separation and detection of simple sugars in standards and experimental samples. The chromatogram includes glucose and fructose standards, as well as cecal digesta samples from germ-free (GF) and conventional (Conv) mice fed inulin- or cellulose-containing diets. Distinct fructose peaks are observed in the fructose standard and in samples from inulin-fed groups (GF and Conv), consistent with the presence of hydrolyzed inulin-derived fructose. No detectable fructose peak was observed in samples from Conv cellulose-fed groups, confirming the absence of fructose in these controls. Retention times of standards were used to identify corresponding peaks in experimental samples.
